# Cosmopolitan Scolytinae: strong common drivers but too many singularities for accurate prediction

**DOI:** 10.1101/2022.05.17.492299

**Authors:** Jean-Claude Grégoire, Hervé Jactel, Jiri Hulcr, Andrea Battisti, Daegan Inward, Françoise Petter, Fabienne Grousset

## Abstract

**Aim:** Many scolytine beetle species have been expanding their range in new territories across geographic barriers, traveling with wood, wood products and plants for planting, sometimes with a high impact on plant health. Here we attempt to quantify the mobility of these cosmopolitan species and to identify the biological drivers of mobility and impact.

**Location:** World

**Major taxa studied:** Coleoptera; Curculionidae; Scolytinae

**Methods:** Mobility was estimated by counting the numbers of landmasses (contiguous pieces of land, surrounded by ocean or sea) colonized by each species. A series of potential drivers (taxonomic tribes; feeding habits; polyphagy; reproductive strategy; host taxa; pheromones and primary attractants) as well as impact on host health were recorded.

**Results:** 163 species were identified, out of 5546 counted in the whole subfamily. Four tribes (Xyleborini; Ipini; Crypturgini; Hylastini) were significantly over-represented, and two others (Corthylini; Hexacolini) were under-represented. 53% of the 163 species are inbreeding, a very significant excess as compared to the whole subfamily (29%). The inbreeders colonized more landmasses than the outbreeders. There is a significant relationship between the number of host families attacked by a species and the number of colonized landmasses. Species restricted to conifers colonized fewer landmasses than hardwood generalists. Species attacking both types of hosts are the most mobile. Most of the invasive species respond to host primary attractants, only one quarter respond to pheromones. All very mobile species respond to primary attractants, and none responds to pheromones. Very mobile species are all associated with a high or moderate impact.

**Main conclusions:** The most mobile species belong for a large part to a limited number of subtribes. They are often inbreeding, polyphagous and respond to primary attractants but do not produce pheromones. However, many species that do not, or only partly, belong to these categories, have established in several landmasses, sometimes with a high impact. For example, the outbreeding *Scolytus multistriatus*, that attacks only 3 host families and produces aggregation pheromones, has established in thirteen landmasses, with a high impact. Therefore, risk prediction needs to assess diversity of species-specific biological traits beyond the few routinely analyzed in literature.

## INTRODUCTION

Very few species are studied in depth before they become noticeable pests. Consequently, most attempts to assess the risk of potentially invasive species rely on limited information. Invasive species assessments now use multiple methodologies ranging from consensus-seeking horizon scans to climate match modeling. However, nearly all these methodologies suffer from one fundamental problem – lack of information about the interactions between a specific species and its potential new environment or hosts.

The typical solution is to take a broader taxonomic perspective and assume that the ecology of a species can be derived from the ecology of related species for which there is more knowledge, or to assume that species within a genus are ecologically similar. The invasive species modeling literature is rich with examples of assessments of genera or even entire families or even guilds (see e. g. Mech et al. 2019; Barwell et al. 2020; Schulz et al. 2021). In this article we demonstrate that even relatively closely related species can differ in their capacity to colonize new territories and in their impact, and we attempt to identify biological and ecological features responsible for these differences.

Some of the most damaging forest pests in the world are bark and ambrosia beetles belonging to the weevil (Curculionidae) subfamily Scolytinae. Global climate change and intense silviculture enabled species such as *Dendroctonus ponderosae* Hopkins and *Ips typographus* L. multiply to epidemic proportions in North America and Europe respectively (Hicke et al. 2016; Grégoire et al. 2015) with a total of 455.7 million m^3^ of pine killed by *D. ponderosae* in British Columbia between 2000 and 2015 (British Columbia Government 2019), and 148 million m^3^ of spruce killed by *I. typographus* between 1950 and 2000 in Europe (Schelhaas et al. 2003), with dramatically increasing damage during the last few years (Hlasny et al. 2021). In addition to these species which are currently spreading within their native continents only, many others have been expanding their territorial range worldwide, especially traveling with international trade. Bark and ambrosia beetles may additionally cause damage as vectors of pathogenic fungi. The redbay ambrosia beetle, *Xyleborus glabratus* Eichhoff, of Asian origin, was first reported in North America in 2002 (Rabaglia et al. 2006). It vectors the fungal symbiont, *Raffaelea lauricola* T.C. Harr., Fraedrich & Aghayeva, causing “laurel wilt”. At least 300 million *Persea borbonia* L. Spreng. (redbay trees) have been killed by laurel wilt in the USA (Hughes et al. 2017), and several other tree species of the Lauraceae family, including avocado (*Persea americana* Mill.) are also affected by the disease. Another example is the introduction^1^ of another Scolytinae, the polyphagous shot hole borer, *Euwallacea fornicatus* Eichhoff (Stouthamer et al. 2017; Smith et al. 2019), which, together with a symbiotic *Fusarium* sp. fungus, attacks a large number of plants, mostly in cultivated settings in its area of origin, Asia, as well as, more recently, in North America (Rabaglia et al. 2006), Israel (Mendel et al. 2012) and South Africa (Paap et al. 2018). The North American species *Dendroctonus valens* LeConte was reported in China at the end of the 1990s and, by 2005, it had spread over 500,000 ha of pine forest in three provinces, killing more than 10 million *Pinus tabuliformis* Carr. (Yan et al. 2005). Other harmful Scolytinae species killing living trees and recently introduced into Europe include the Asian ambrosia beetle, *Xylosandrus crassiusculus* (Motschulsky), the black twig borer, *X. compactus* (Eichhoff), and the walnut twig beetle, *Pityophthorus juglandis* Blackman. This latter species vectors the pathogenic fungus *Geosmithia morbida* Kolarík, Freeland, Utley & Tisserat, causing thousand cankers disease of walnuts, *Juglans* spp. (EPPO 2015; Seybold 2019).

The observed spread of many species continues. At the same time, dozens of bark- and ambrosia beetle species have been introduced into non-native regions without any detectable impact. Most bark beetle “tramp species” are harmless.

So far, at least 163 species out of the ∼6,000 described scolytine species (Hulcr et al. 2015) have established outside of their native areas (Tables 1. and S-1). The remaining ninety-eight percent of scolytine species are thus still potentially able to colonize new territories, and their potential impact is still mostly unknown.

**Table 1.**
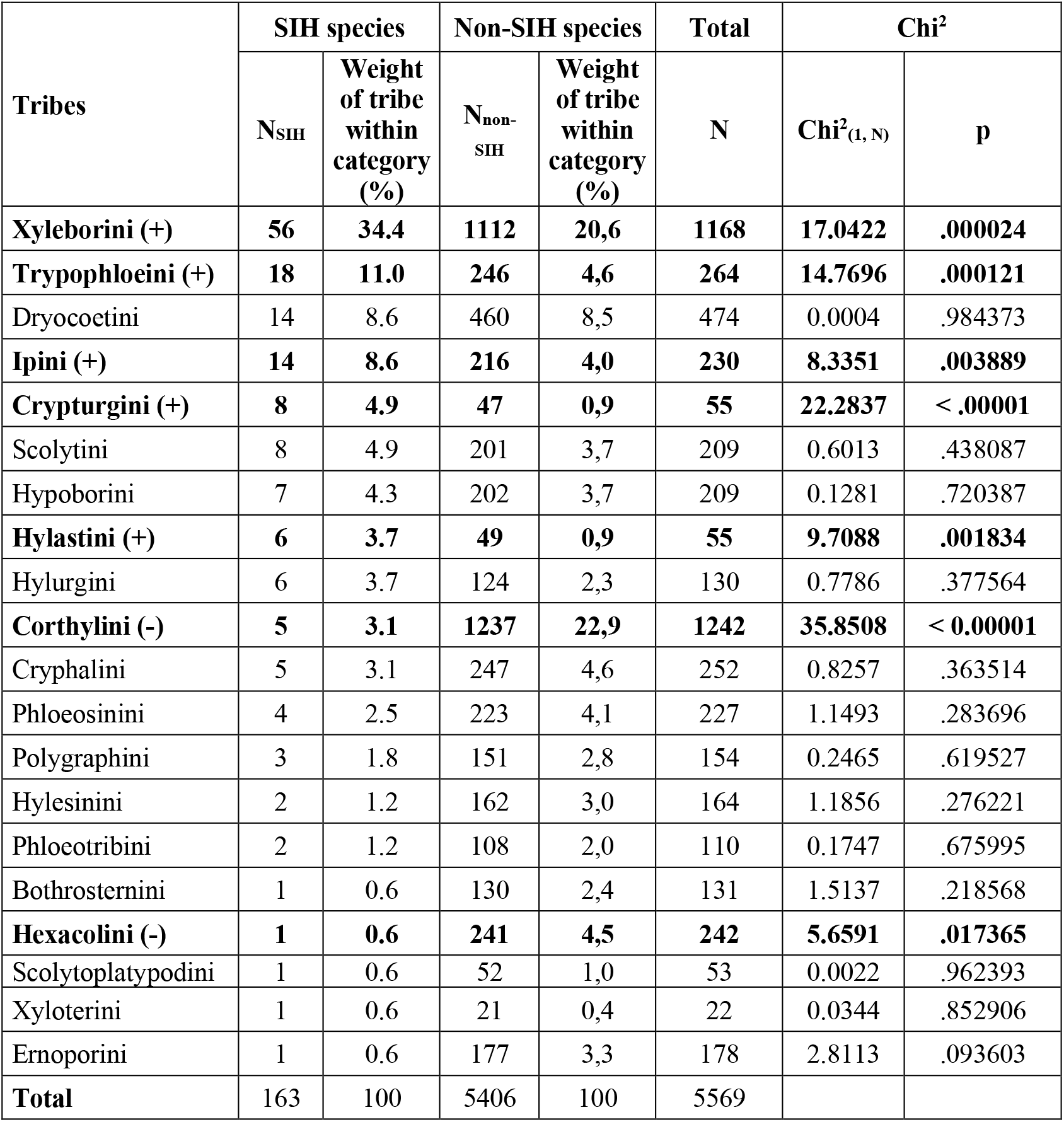
Tribes represented among the Scolytinae with an invasion history (SIH). Tribes over-represented among the invasive Scolytinae are in bold, followed by (+); tribes under-represented are in bold, followed by (-). World figures taken from Hulcr et al. (2015)), except for the Trypophloeini, Cryphalini, Corthylini and Ernoporini, for which the revision by Johnson et al. (2020) was used. The number of non-SIH species is calculated by substracting the number of SIH in a tribe from the total number of species in the tribe.

Other species that have not spread to date, and which are not recognized as harmful, might start expanding their range, benefiting from the trade of new commodities or from commercial movements along new routes. These beetles, alone or together with pathogens, may also colonize new hosts that may prove to be more susceptible than their native hosts, or form new associations with local pathogens as suggested by Rassati et al. (2019a). For both known and unrecognized spreading species, the possibility that they can be successfully introduced into new areas, and their subsequent potential economic or environmental impact are two major components of phytosanitary risk.

“Horizontal” regulations globally addressing the host plants of non-native pests are locally implemented. For example, all non-European Scolytinae attacking conifers are targeted in the European Union by phytosanitary requirements applying to the importation of coniferous wood^2^ (EU 2019) but equivalent requirements do not exist for the trade of non-coniferous wood. A recent EPPO study focused on twenty-six representative Scolytinae and Platypodinae ambrosia- and bark beetle species associated with non-coniferous wood (EPPO 2020; Grousset et al. 2020). Sixteen life-history traits and other factors were qualitatively weighed with expert knowledge against invasion success. Inbreeding, polyphagy (number of host families) and the lack of aggregation pheromones were common features of species with a successful introduction history. Association with pathogenic fungi, the use of aggregation pheromones, and the capacity to attack and kill new host species were identified as factors contributing to high impact. One of the important conclusions of this EPPO study was that traits related to species with a past invasion history had a strong influence on invasion risks. However, it was found that the main factors that are driving successful establishment and impact vary from species to species and are not always fully identified. One important recommendation of this study was that horizontal phytosanitary measures similar to those for conifer wood better address the risk than regulation of individual species. In another recent study, Lantschner et al. (2020) similarly reviewed 123 Scolytinae species with a history of invasion, focusing on biological characteristics (feeding habits and mating systems), cumulative trade between world regions, size of source species pools, forest area and climatic matching between the invaded and source regions. They identified sib-mating as a major factor favoring the movement of Scolytinae species into new territories, but also found that a non-biological trait, cumulative trade between world regions is a primary driver of scolytine invasion.

At a broader taxonomic scale, Mech et al. (2019) and Schulz et al. (2021) focused on the impact of non-native herbivorous insects established in North America. They found that the evolutionary proximity between the native and novel host plants, life history traits of the novel hosts, and the presence of native close congeners with a long-term association with the novel host were better predictors of impact than were traits of the invading insects themselves.

Presence in at least one new area was used as a criterion to select the species analysed in these studies. Here we attempt to quantify further the introduction capacity of 163 Scolytinae species known for having moved to new territories.

## METHODOLOGY

A dataset of Scolytinae species known to have spread beyond geographical barriers (across seas or oceans in this study) was constructed (Table S-1), including any species distributed across at least one barrier (hereafter designated as “*Scolytinae with an invasion history*” - SIH), irrespective of its area of origin which is often difficult to establish (see e. g. Lin et al. 2021). The list includes all the Scolytinae species from the EPPO study (EPPO 2020; Grousset et al. 2020), as well as the species introduced in North America, New Zealand, and Europe, listed respectively by Haack (2001, 2006), Brockerhoff et al. (2006) and Kirkendall and Faccoli (2010). This initial set was expanded using information mostly from Wood and Bright (1992), Atkinson (2021), Lantschner et al. (2020), and from other publications (full list of references in Table S-1). The dataset was completed in December 2020 and, therefore, does not include several important studies (in particular Bright 2021) published after this date. Several biological features were considered. Fungus farming, although considered in previous studies (EPPO 2020; Grousset et al. 2020), was not included as a separate factor in this analysis because it applies to all ambrosia beetles, while among bark beetles, mycetophagy is still poorly documented. Similarly, the association with pathogens was not considered as a predictor since, in addition to previously known species, species so far harmless on their native hosts (e. g. *R. lauricola, G. morbida*) become pathogenic when their vectors colonize new host trees. Besides, scolytines species considered as harmless are sometimes found associated with aggressive pathogens (Wingfield and Gibbs 1991), making pathogens a dubious predictor of impact. Climatic requirements, dispersal capacity and voltinism were also not considered, because of the wide knowledge gap regarding these potential drivers (but see EPPO 2020 and Grousset et al. 2020). The counting of colonized landmasses served as a proxy to estimate long-distance dispersal.

### Feeding regimes

We retained the following general categories (Kirkendall et al. 2015): *phloeophagy* (feeding in inner bark; this category corresponds to the bark beetles *stricto sensu*); *xylomycetophagy* (fungus farming; this category corresponds to the ambrosia beetles, which live in the xylem of woody plants, where they cultivate symbiotic fungi on which they feed); *spermatophagy* (feeding in seeds) and *herbiphagy* (feeding in non-woody plants).

### Inbreeding

Unless specified in Table S-1, the information comes from Kirkendall et al. (2015).

### Polyphagy

Polyphagy was measured, as in EPPO (2020) and Grousset et al. (2020), by the number of host-plant families colonized. Unless specified otherwise in Table S-1, host-plant data come from Atkinson (2021) or Wood and Bright (1992a).

### Aggregation pheromones (categories: 0/1/2)

The source for this field is El-Sayed (2018) unless specified otherwise. Three categories were considered: 0 (no pheromone known for the genus); 1 (pheromones known for other species in the genus); 2: (pheromones known for the species).

### Primary attractants (0/1/2)

Unless specified by a footnote, the information regarding primary attractants (e. g. ethanol and/or alpha-pinene, emitted by the host or by other organisms within the host) comes from Atkinson (2021). The three categories considered are the same as for pheromones.

### Host plants: conifers vs. non-conifers (1/2/3)

Three categories were considered: 1 (species attacking only conifers); 2 (species attacking only non-conifers); 3 (species attacking both conifers and non-conifers).

### Impact on plant health (0/1/2)

Three categories: 0 (no impact); 1 (moderate impact); 2 (substantial impact). Criteria for damage by spermatophages: massive colonisation of fruits and/or impact on regeneration.

### Landmasses

We use the term *landmass* to define a contiguous piece of land (a continent or an island, irrespective of its size) surrounded by ocean or sea. This approach admittedly creates large biases. Even if a continent is very large, we consider it as a single landmass. The movements of a species within a landmass are not considered because they are often incompletely documented. However, continents that are not fully separated by oceans (North, Central and South America; Europe, Asia, and Africa) are considered as distinct landmasses because of the distances and ecoclimatic differences between them. Some archipelagos (e. g. Cape Verde, Fiji, Galápagos, Hawaii, Micronesia) were considered each as one unit. Islands comprising several countries (e. g. Republic of Ireland + Northern Ireland; Haiti + Dominican Republic) were considered as single units. The size of the geographic barriers between landmasses and of the landmasses themselves has not been considered. Great Britain and the European mainland would thus be considered as separate landmasses although the Channel that separates them is locally less than 35 km broad. On the other hand, South America, which is more than 7000 km long, is considered as a single landmass. Despite these many inconsistencies, we believe that this approach provides a useful, if probably conservative, metric to consider pest mobility. Table S-1 provides a listing and a counting of the discrete landmasses occupied by each species. The acronyms used to designate the different landmasses are listed in Table S-2. When possible, ISO alpha-3 codes (https://www.iso.org/obp/ui/#search) were used. Codes for locations absent from this list because they refer to intra-national territories (e. g. an island belonging to a larger country) were taken from the International Working Group on Taxonomic Databases For Plant Sciences (TDWG) (https://github.com/tdwg/wgsrpd) or were created for the purpose of this analysis.

### Statistical analyses

#### Weight of the different tribes among the SIH; feeding regimes *vs*. reproductive strategies

2 × 2 Chi-Square tests were used, with Yate’s correction for continuity for expected values inferior to 5.

#### Multivariate analyses on impact

A factorial discriminant analysis (FDA) was performed as a supervised classification method to discriminate between three categories of beetle species *a priori* classified, as in the Methodology and in Table S1, according to their level of damage (impact), as having no impact (0), moderat impact (1) or substantial impact (2), using ecological characteristic as predictor variables (Table S-1). The data set consisted of 163 species characterized by one quantitative functional trait, polyphagy, expressed as the number of known host plant species, and five qualitative functional traits transformed into dummy variables, namely whether bark beetle species exhibited the following characteristics: xylomycetophagy (ambrosia beetles), inbreeding, using aggregation pheromones, using primary attractants, and host specialization (“specialists”: attacking either conifers or non-conifers; “generalists”: attacking both).

#### Covariance analyses on mobility

A Spearman correlation analysis was performed between the number of colonized land masses and the functional traits of the 163 scolytine species. Two variables were identified as significantly correlated with beetle cosmopolitanism, one quantitative, the degree of polyphagy (expressed in terms of number of known host plant families) and one qualitative, the use (or not) of primary attractants for host plant colonization. We then used an analysis of covariance (Ancova, with and without interaction) to assess the magnitude of the effects of these two factors. All statistical analyses were made with XLSTAT.

## RESULTS

### 1. Scolytinae with an invasion history - overall features

#### 1.1. Representation of the different tribes of Scolytinae

Five tribes, the Xyleborini, Trypophloeini, Ipini, Crypturgini and Hylastini are significantly more frequent among the invasive Scolytinae than among the Scolytinae as a whole. Two tribes, the Corthylini and Hexacolini are significantly less frequent (Table 1).

The small tribes Amphiscolytini (1 sp.), Cactopinini (21), Carphodicticini (5), Hyorrhynchini (19) and Phrixosomatini (25) are absent from the list, as well as the larger tribes Diamerini (132), Micracidini (298) and Xyloctonini (78).

##### Feeding regimes

Amongst the 163 SIH species, 79 (48.5 %) are phloeophagous, 60 (36.8 %) are xylomycetophagous, twelve (7.4 %) are herbiphagous and twelve are spermatophagous. A majority (82.3%) of the phloeophages among the SIH are outbreeding, whilst a majority of the xylomycetophages (93.3%) and of the spermatophages (83.3%) are inbreeding. The mating habits of the herbiphages are equally balanced (Table 2).

**Table 2.** Feeding regimes of the Scolytinae with an invasion history

| Feeding regime | Outbreeding |  |  | Inbreeding |  |  | Total |  | Chi<sup>2</sup><sub>1</sub> |  |
| --- | --- | --- | --- | --- | --- | --- | --- | --- | --- | --- |
|  | N | % of total | % of regime | N | % of total | % of regime | N | % of total | Chi<sup>2</sup><sub>1</sub> | p |
| Xylomycetophagy (+) | 4 | 2.5 | 6.7 | 56 | 34.4 | 93.3 | 60 | 36.8 | 60.9222 | < 0.00001 |
| Phloeophagy (-) | 65 | 39.9 | 82.3 | 14 | 8.6 | 17.7 | 79 | 48.5 | 78.3002 | < 0.00001 |
| Herbiphagy | 5 | 3.1 | 41.7 | 7 | 4.3 | 58.3 | 12 | 7.4 | 0.128 | .720506 |
| Spermatophagy (-) | 2 | 1.2 | 16.7 | 10 | 6.1 | 83.3 | 12 | 7.4 | 4.6719 | .03066 |
| Total | 76 | 46.6 |  | 87 | 53.4 |  | 163 |  |  |  |

### 2. Biological features influencing risks of introduction and impact

#### 2.1. Mating strategy

Among the 163 species in our study, 87 (53.4%) are inbreeding (Table 3). Nearly all females leaving the tree are thus already fertilised and can create a new colony on their own. In theory, the Allee population threshold (the minimal number of individuals below which a population cannot grow) for such species could be one single female. This proportion of inbreeding species is significantly larger than that (27.8%) of the non-SIH inbreeders in the world (1544 species - Kirkendall et al. 2015) among the known species belonging to tribes with SIH species (5569 species - Hulcr et al. 2015; Johnson et al. 2020): Chi^2^_(1; N=5569)_ = 47.42; p<0.00001. The Xyleborini and Trypophloeini, overrepresented in Table 1, are all inbreeding, and the underrepresented Corthylini and Hexacolini are all outbreeding. However, the overrepresented Crypturgini and Hylastini are all outbreeding (Table 3).

**Table 3.**
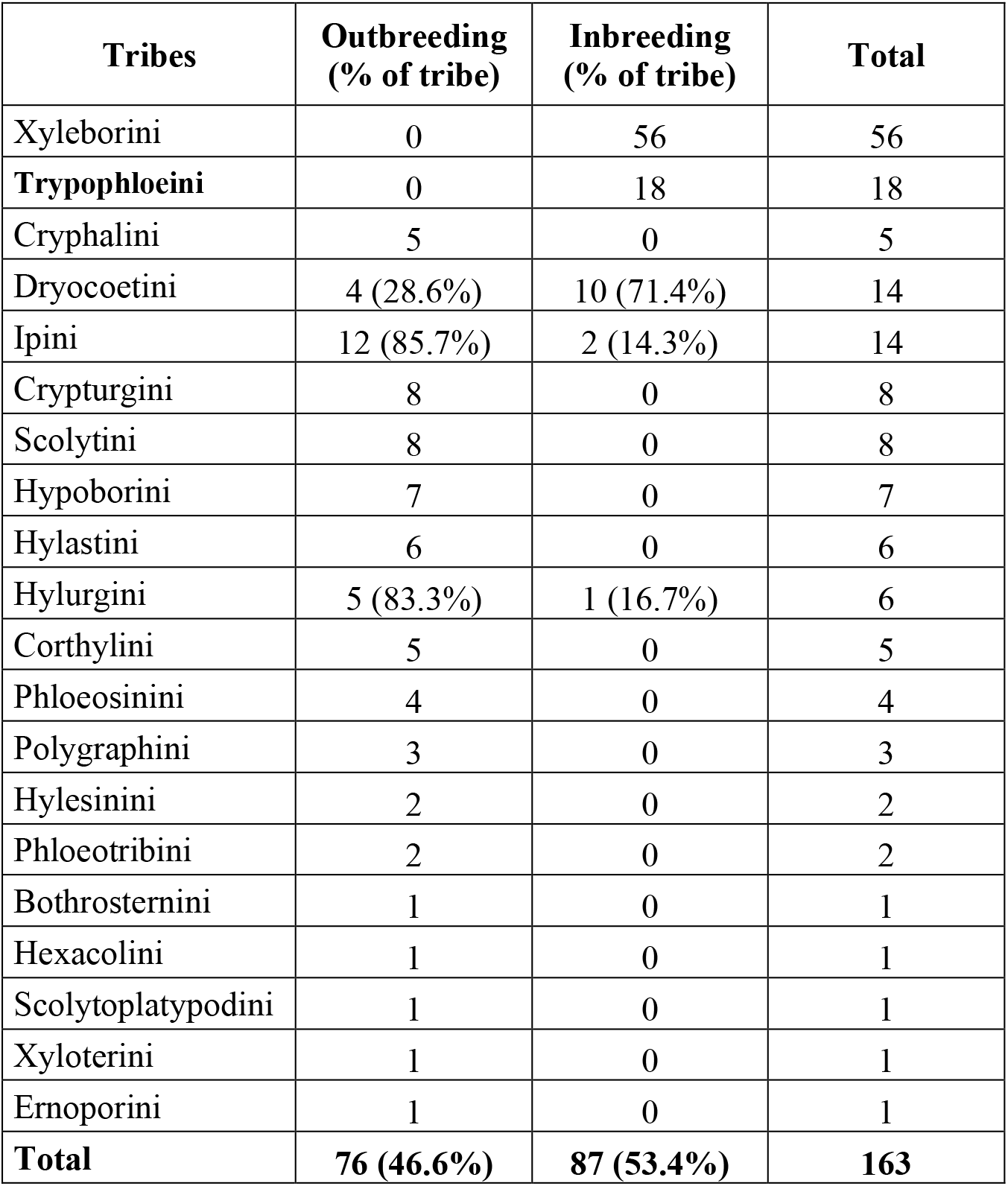
Mating strategies of the Scolytinae tribes with an invasion history

Inbreeders are also often haplodiploid. Unfertilised females parthenogenetically produce haploid males, and then mate with their son (Jordal et al., 2001, and references therein). This further facilitates colonisation as females do not even have to be fertilised before dispersal and finding a host. For example, all the Xyleborini and most of the *Coccotrypes* spp. are haplodiploid (Jordal et al. 2001; EPPO 2020; Grousset 2020).

Among the supposedly outbreeding species that crossed a geographic barrier, *Orthotomicus erosus* (Wollaston) (Mendel 1983) and *Tomicus piniperda* (Linnaeus) (Janin and Lieutier 1988) show a proportion of females already mated upon emergence, possibly with a sibling, or mated during maturation feeding on twigs or during overwintering at the base of trees previous to colonising a new host. Similarly, *Hylurgus ligniperda* (Fabricius) (Fabre and Carle 1975) and *Ips grandicollis* (Eichhoff) (Witanachchi 1980) have been observed to mate prior to emergence. As in the inbreeding species *stricto sensu*, these early mated females may be able to start a new colony alone. Wilkinson (1964) showed that single *I. grandicollis* females induced to oviposit on pine logs produced a progeny. However, species with no invasive history are also capable of early mating. Lissemore (1997) found that 3 out of 8 pre-emergent, overwintering *Ips pini* (Say) females collected in the spring in the litter around attacked trees were fertilized and able to start a new gallery alone. The North American species *Ips pini* has never expanded outside of its range, where it is widely distributed (Atkinson 2021). Similarly, Bleiker et al. (2013), examining 1510 emergent female *Dendroctonus ponderosae* Hopkins from two different locations in Alberta, found 3-5% of preemergent matings. Overall, the inbreeding (*stricto sensu*) SIH colonized a much larger set of landmasses than the outbreeding species (Figure 1). Strikingly, with the exception of *Hypocryphalus mangiferae* (Stebbing) (17 landmasses), all the species colonising the larger numbers of landmasses are inbreeding.

**Figure 1.**
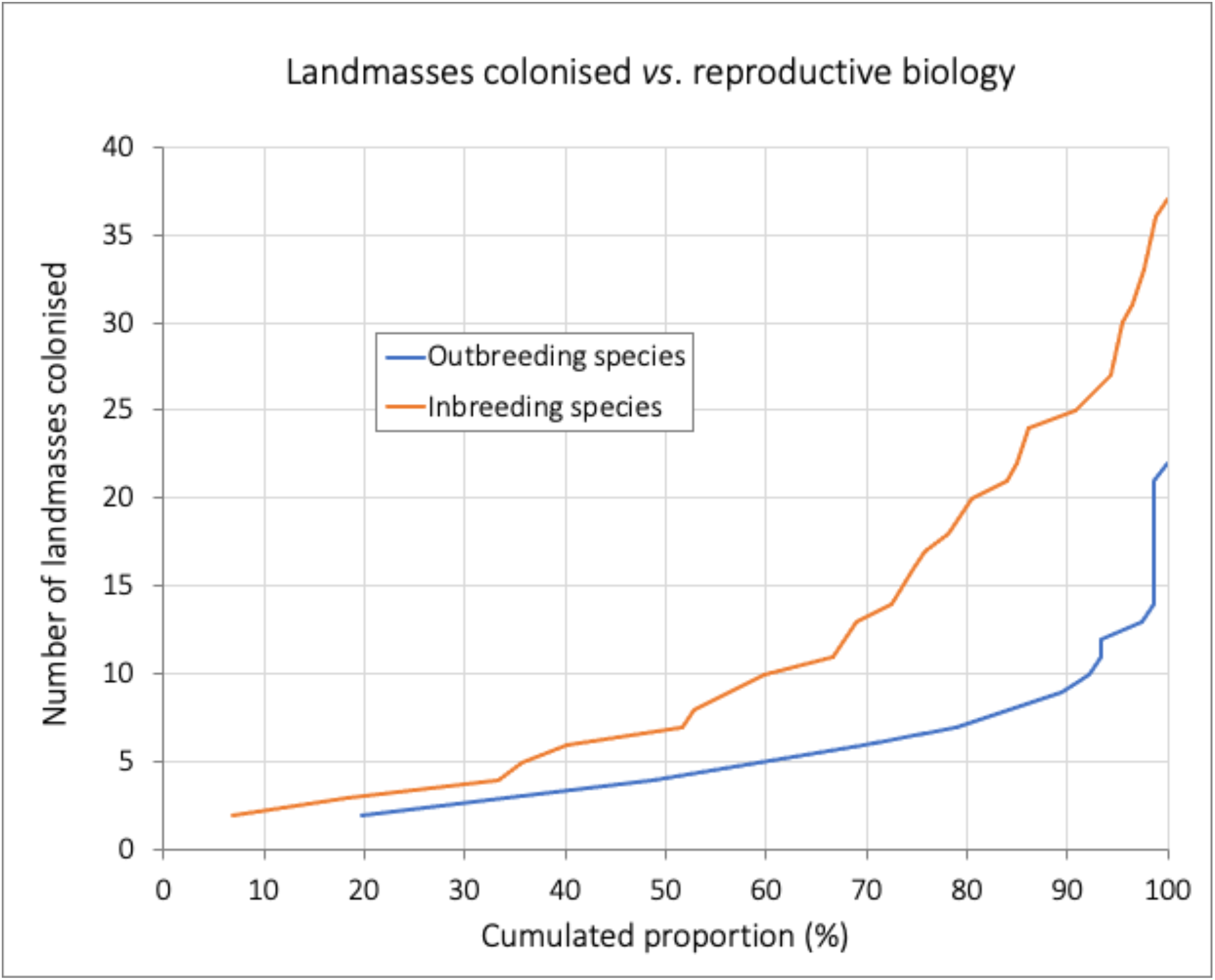
Cumulative proportion of landmasses colonized by either outbreeding or inbreeding species among the Scolytinae with an invasion history

#### 2.2. Host nature and condition

Many different relationships to the hosts are observed among bark- and ambrosia beetles, making it difficult to predict the risks associated with new insect-host associations, or even the long-term risks associated with long-standing associations. Bark- and ambrosia beetle species attack a wide range of trees, from apparently healthy individuals to dead and even decaying ones (Raffa et al., 2015; Hulcr et al., 2017). Other SIH species colonize a wide range of plant parts and therefore commodities in trade, including seeds, fine twigs and roots (Kirkendall et al. 2015, and see section 1.2). The nature and condition of the host allow a certain level of prediction regarding the entry, establishment, and impact of a particular species or, after an event has occurred, provide clues for retrospective scenarios.

##### Entry

Xylophagous and xylomycetophagous species living in the sapwood are protected from mechanical damage and, when the wood has not been dried, from desiccation. Many phloeophagous bark beetles (e. g. *H. ligniperda*) and xylomycetophagous ambrosia beetles (e. g. *Xylosandrus germanus*) (Blandford) have travelled in wood packaging material, or in wood or wood product shipments. Plants for planting provide another pathway for species living in the stems of living hosts, such as *Xylosandrus compactus* (Eichhoff). *Coccotrypes dactyliperda* Fabricius, which live in dates, is likely to have spread around the world in commercial shipments. *C. rhizophorae* (Hopkins), which specifically lives in the propagules of the red mangrove, *Rhizophora mangle* Linnaeus, might have moved from Asia where it originates to North America in host propagules floating long distance across the ocean (Atkinson & Peck 1994).

##### Establishment and impact

Species capable of attacking living trees are more likely to find suitable hosts in the locations of entry. Hulcr et al. (2017) proposed to search for ambrosia beetle-fungus associations colonising live trees in their native habitats to identify future exotic tree-killing pests. Living trees, however, can vary in vigour and resistance to pests. Often, apparently healthy trees have been previously exposed to various forms of stress factors, including flooding, drought, wind break, snow break, freezing, ozone exposure, graft incompatibility, site and stand conditions, nutrients supply disorders, diseases, or animal pest damage (Ranger et al., 2010; Hulcr and Stelinski, 2017; Ploetz et al., 2013; Brune, 2016), and this generally makes them more vulnerable to beetle attacks. Thirty-five SIH species may kill stressed hosts; twenty-one species out of 163 are able to kill apparently healthy, living trees (Table S1B).

Importantly, impact in a new area cannot always be predicted from the relationship of a beetle-fungus association with its native host trees. *X. glabratus* and its symbiont *R. lauricola* colonize stressed or injured Lauraceae all over the world. Whilst they exert little noticeable damage in their native areas, they massively kill *P. borbonia* in the USA because of the hypersensitive response of the New World Lauraceae and the changes in behaviour they induce in the beetles (Hulcr et al. 2017; Martini et al. 2017). *Anisandrus dispar* (Fabricius), which attacks weakened or dead trees in Europe is an important pest of young chestnut trees stressed by excess water or late frost in north-western USA and western Canada (Kühnholz et al. 2001). Similarly, *D. valens*, which usually settles on the stumps of freshly cut pines or more rarely establishes in low numbers on stressed pines in North America, killed millions of *Pinus tabuliformis* since its introduction in China during the late 1990s (Yan et al. 2005). The causes of this increased aggressivity in China are unclear but have been related to exceptionally dry years following introduction (the outbreak subsided after the drought) and, possibly to some degree, to the association with a new, naïve host, with more aggressive strains of symbiotic fungi (Sun et al. 2013). Sometimes, even in their native range, species usually restricted to dead or dying hosts start attacking apparently healthy trees.

*Trypodendron domesticum* (Linnaeus) and *T. signatum* (Fabricius) started infesting thousands of standing, live beech *Fagus sylvatica* L. in Belgium in the early 2000s, in connection with exceptional early frosts (La Spina et al. 2013). In Canada, *T. retusum* (LeConte) which is usually restricted to wind-broken or weakened trees was observed to attack apparently healthy aspen, *Populus tremuloides* Michaux (Kühnholz et al. 2001).

Overall, the capacity to colonize living hosts appears to favor establishment. In our dataset, species with a recorded impact on their hosts colonized the larger numbers of landmasses (Figure 2).

**Figure 2.**
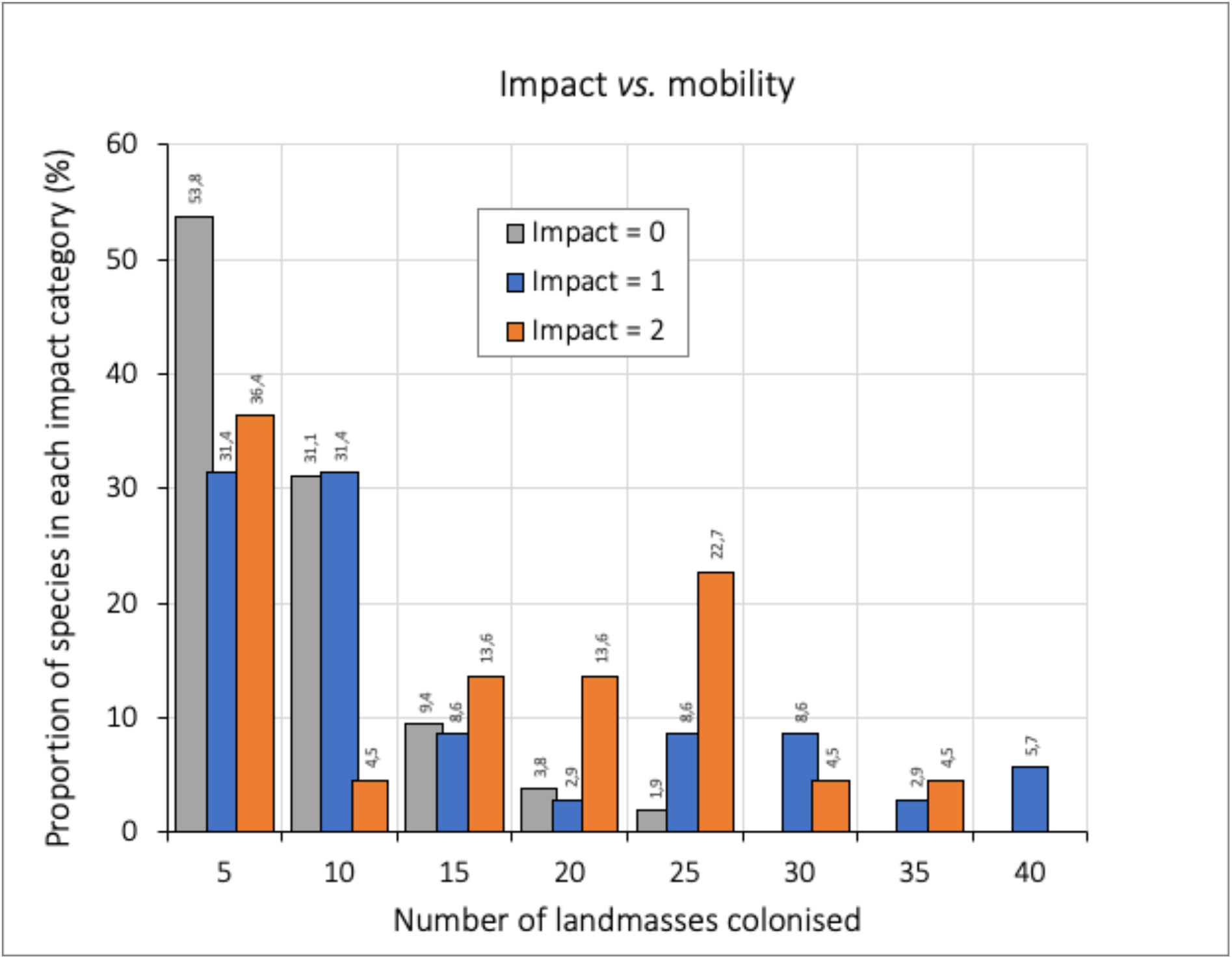
Impact versus mobility among scolytines with an invasion history

#### 2.3. Host specificity

Polyphagy and the ability to attack new hosts in new locations are advantageous for entry, establishment (higher probability of finding a suitable host) and impact (EPPO, 2020).

##### Polyphagy

Bark beetles usually have a narrow host range and are often monophagous (all hosts belong to the same genus) or oligophagous (all hosts selected within one family). Ambrosia beetles often have a broader range of hosts, as their host is mainly a substrate for the fungi they grow and feed on (Beaver 1979; Jordal et al. 2000; Seybold et al. 2016). Many species specialize in either conifers or non-conifers, although some exceptionally polyphagous species attacks both.

Among the 36 species in Table S-1 attacking only conifers, 33 species attack only one family, and two species attack two families. The Scolytinae attacking only non-conifers or attacking both non-conifers and conifers have a much wider and diverse range of host trees. Conifer specialists colonize fewer landmasses (median: 5) than non-conifer specialists (median: 6) and species attacking both types of hosts (median: 9): Figure 3.

**Figure 3.**
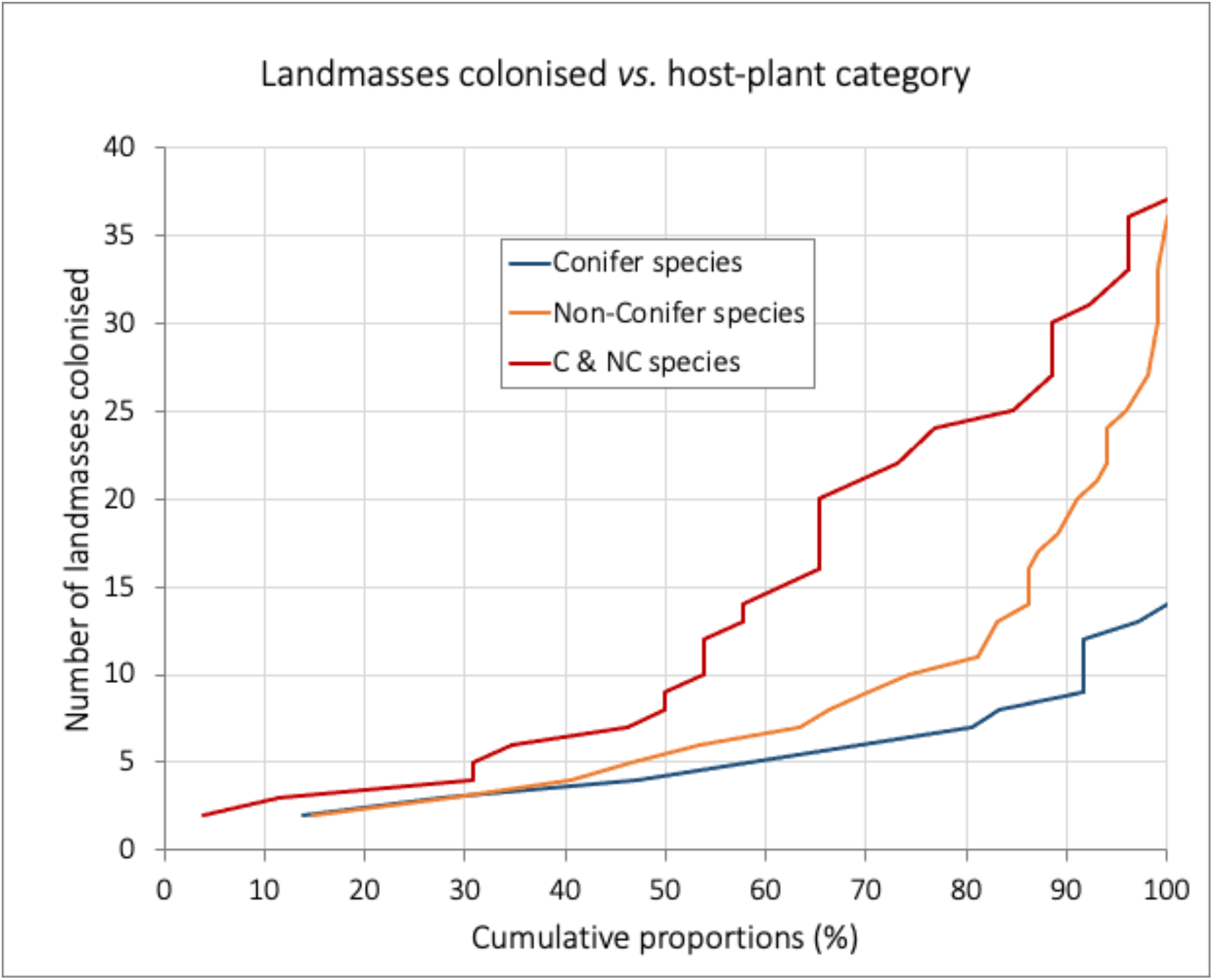
Host-plant category (conifer vs. non-conifer) influences the number of landmasses colonized by Scolytinae with an invasion history

The genus *Hypothenemus*, representing 11% of the 163 species in the list, includes the most polyphagous species in the list with *H. eruditus*, reported from 65 plant families, and *H. crudiae* and *H. seriatus*, each reported from 57 plant families. These species are reported from 37, 21 and 22 landmasses, respectively.

There is no direct relationship between polyphagy and impact. Some less polyphagous ambrosia beetles have a substantial impact in newly invaded territories, as illustrated by *X. glabratus* (4 host-plant families) after its introduction in the USA). On the contrary, very polyphagous species may cause limited damage in new areas as well as in their native range. *Hypothenemus eruditus*, (65 host families), which usually colonizes dead hosts, is usually considered harmless (Kambestad et al. 2017).

##### New hosts

Many scolytines, even some not known as polyphagous, have been recorded on new host species when introduced into new areas (EPPO 2020; Grousset et al., 2020).

Encounters with new hosts do not always result in damage but are an important component of the potential impact. There are striking example of encounters with new very susceptible hosts, leading to extensive damage, such as *X. glabratus* on *Persea borbonia* in the USA (EPPO, 2020), or *D. valens* on *P. tabuliformis* in China (Yan et al. 2005).

#### 2.4. Aggregation pheromones and primary attractants

##### Aggregation pheromones

The need for mass-attacks can be unfavorable to establishment, but mass attacks once the species is established and the epidemic threshold is reached can result in higher impact (EPPO, 2020). Some bark or ambrosia beetles use aggregation pheromones to mass-attack standing hosts and overcome their defences (D.L. Wood 1982). The mass-colonisation of undefended, fallen trees is more likely the result of collective foraging, also mediated by aggregation pheromones (Toffin et al. 2018). As large numbers of individuals are required for a mass-attacking species to colonize a new tree, the Allee threshold is necessarily high, making establishment in a new area more difficult. On the contrary, solitary colonizers (e. g. *Hypothenemus* spp.; *Xylosandrus* spp.) have displayed high success in establishment (see section 2.1).

Once the species is established and the epidemic threshold is reached, aggregation pheromones facilitate mass attacks on live trees, which results in higher impact. Such mass attacks are known among the SIH species, i.e. for *Orthotomicus erosus, Gnathotrichus materiarius* (Fitch), *Ips calligraphus* (Germar), *I. cembrae* (Herr), *I. grandicollis, Pityogenes bidentatus* (Herbst), *P. calcaratus* (Eichhoff), *P. chalcographus* (L.), *Pityokteines curvidens* (Germar), *Pityophthorus juglandis, Polygraphus poligraphus* (L.), *P. proximus* Blanford, *P. rufipennis* (Kirby), *Scolytus amygdali* Guerin-Meneville, *S. multistriatus* (Marsham), *T. domesticum* and many others.

##### Primary attractants

Physiologically stressed trees emit a range of volatile compounds, such as ethanol, which attract many bark and ambrosia beetles colonising weakened hosts (Byers 1992; Miller and Rabaglia 2009; Ranger et al. 2010; Rassati et al. 2019b). Monoterpenes emitted by conifers also serve as clues for conifer-inhabiting species (Byers 1992) but reduce the response of species attacking non-conifers to ethanol or other primary attractants (Ranger et al. 2011). The coffee berry borer, *Hypothenemus hampei* is attracted to ripe coffee berries by conophthorin and chalcogran but deterred by conifer monoterpenes (Jaramillo et al. 2013). Beetle response to primary attractants can be extremely accurate. In South Africa, Tribe (1992) showed that adults of the European species *Hylastes angustatus* (Herbst) and *Hylurgus ligniperda* were capable of finding *Pinus radiata* logs buried horizontally under 40 cm of soil. This accuracy is perhaps one component of the invasive success of these two species. However, working with native secondary species in Canada, Saint-Germain et al. (2007) showed that primary attractants allow bark beetles to locate a patch inhabited by susceptible hosts but that, at closer range, host selection is governed by different processes, including random landing.

Because they are not very specific (e. g. ethanol is produced by tissue fermentation of both conifers and non-conifers, and monoterpenes such as alpha-pinene are produced by most conifers), primary attractant can particularly facilitate host location and thus establishment among polyphagous species. 94 SIH species out of 163 are known to respond to primary attractants, and an additional 47 are likely to use these chemical clues as well.

Twenty species are not known to respond to primary attractants and do not produce pheromones either: five *Aphanarthrum* spp.; *Dendroctonus micans*; *Dryoxylon onoharaense*; *Kissophagus hederae*; six *Liparthrum* spp.; *Microborus boops*; two *Microperus* spp; *Pagiocerus frontalis*; *Scolytoplatypus tycon*; *Thamnurgus characiae*.

#### 2.5. Multivariate analyses

##### Impact

The factorial discriminant analysis showed significant effects of functional traits on impact (Wilks’ lambda test, P<0.0001). The separation between the three impact levels was mainly explained by the FDA canonical function F1 (percentage variance explained 81.8%, P < 0.0001; while F2 explained 18.2%, P = 0.09). F1 was mainly driven by the degree of polyphagy (P=0.001), use of aggregation pheromones (P=0.002), host specialization (P=0.004), and to a lesser extent, use of primary attractants (P=0.089). The confusion matrix (Table 4) showed 100% correct classification for the category of non-damaging beetles (no impact; 107 species) The beetle species with no impact were characterized by a low degree of polyphagy, lack of aggregation pheromone, host specialization on broadleaves or conifers, and non-use of primary attractants. Only 11.4% of scolytine species with moderate impact and 9.5% with substantial impact were correctly classified, the other species of these categories being mainly mis-classified as non-damaging. However, it should be noted that four *Euwallacea* species combined traits of polyphagy and lack of host specialization, using of aggregation pheromone and primary attractant: *E. piceus, E. interjectus, E. similis*, and *E. validus* and they all had a significant impact.

**Table 4.** Confusion matrix for the Factorial Discriminant Analysis (FDA) of the three categories of impact by the 163 beetle species studied The complete list of well-classified and mis-classified species is available as supplementary material (Table S-3).

| *a priori* \ *a posteriori* | No impact | Low impact | Substantial impact | Total | % correct |
| --- | --- | --- | --- | --- | --- |
| **No impact** | 107 | 0 | 0 | 107 | 100% |
| **Moderat impact** | 29 | 4 | 2 | 35 | 11.4% |
| **Substantial impact** | 15 | 4 | 2 | 21 | 9.5% |
| **Total** | 151 | 8 | 4 | 163 | 69.3% |

##### Mobility

The Ancova analysis showed a significant effect of the degree of polyphagy (P<0.0001) and use of primary attractant (P=0.023) on the number of landmasses colonized, but the interaction of these two factors was not significant (P=0.58), with an overall determination coefficient of R^2^=0.41. Beetle species not using primary attractants (n=22) colonized significantly fewer land masses (3.5 ± 0.4, mean ± standard error) than those (n=141) attracted by the host plant (9.6 ± 0.7). The number of colonized land masses increased with the degree of polyphagy (number of known host plant species), by the same magnitude for the two categories of beetle species (using or not primary attractants, Figure 4).

**Figure 4.**
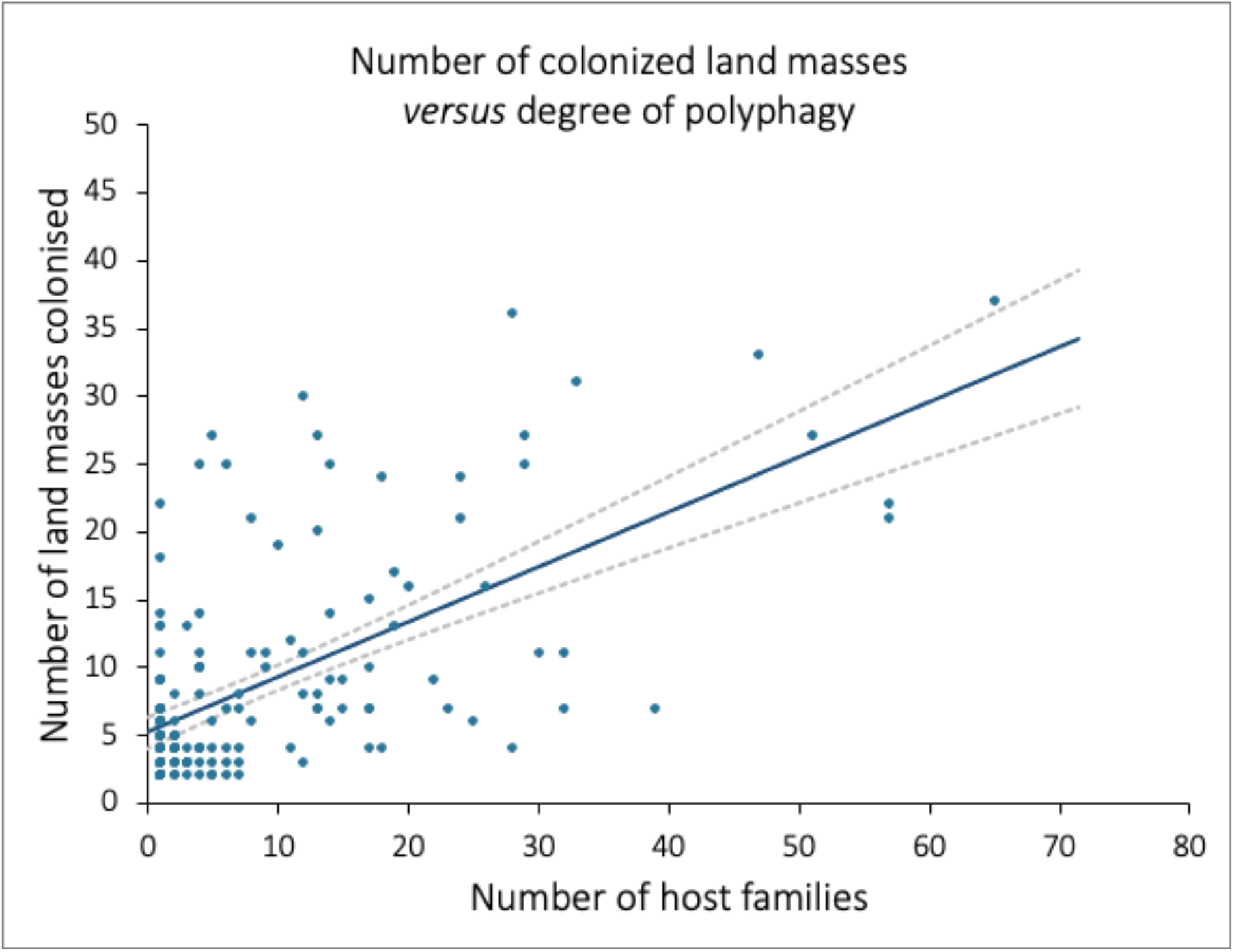
Number of colonized land masses *versus* degree of polyphagy (number of host plants) for the 163 scolytine species studied (independently of their use of primary attractants). Dashed lines represent the confidence interval of the linear regression line

## DISCUSSION AND CONCLUSION

Throughout this review, several biological traits, particularly inbreeding and polyphagy, appear correlated with higher introduction and impact in new areas. However, as with the results obtained in EPPO (2020) and Grousset et al. (2020) for a narrower range of species, none of these traits, alone or combined, explains the success of all the SIH species, and there are obvious outliers. For example, the over-represented tribes Crypturgini and Hylastini (Table 1) are outbreeders. The moderately polyphagous *X. glabratus* (4 host families) has a much higher impact than *H. eruditus* (65 host families). More generally, 59 SIH species attack hosts in only one plant family, suggesting many exceptions to the influence of polyphagy on introduction. Whilst aggregation pheromones do not appear to favor establishment (Figure 7), there is the exception of *E. fornicatus*.

To summarise, some of the identified drivers are widespread among SIH species, but none are shared by the whole group, making it difficult to characterize univocally the potentially successful invaders among the bark and ambrosia beetles of the world. In addition, the non-biological risk factors as identified in EPPO (2020) and Lantschner et al. (2020) also play an important role. As concluded in EPPO (2020), the main factors that are driving successful establishment and impact vary from species to species and are not always fully identified. Still, one single feature common to most of the SIH species has been implicitly identified in this study on species crossing geographical barriers: their capacity to travel by trade, either on wood commodities and wood packaging material, or on plants for planting, or on fruits, depending on the species. The major conclusion of the present study is thus that, because of the lack of drivers that could allow for robust predictions regarding the invasive potential of any scolytine species, it is safer to consider the establishment of horizontal measures for trade of commodities.

## AUTHOR CONTRIBUTIONS

J.-C.G. conceived the study; J.-C.G. and H.J analysed and interpreted the data; J.-C.G. compiled the data, produced the figures and wrote the initial draft; J.H. particularly supervised the taxonomy issues ; F.G. and F.P. particularly supervised the plant health regulatory issues; all authors supervised the ecology issues and contributed substantially to the revision of the initial draft.

## Supporting information

Table S1: Main characteristics of the 163 Scolytinae with an invasion history.

Table S2: Landmasses (islands and continents).

Table S3: Well classified and mis-classified species identified by the Factorial Discriminant Analysis (FDA) of the three categories of impact by the 163 beetle species studied.

## Footnotes

1 In this context, ‘the entry of a pest resulting in its establishment’, following the terminology of the Glossary of Phytosanitary Terms of the International Plant Protection Convention (FAO 2019).

2 ‘Commodities such as round wood, sawn wood, wood chips and wood residue, with or without bark, excluding wood packaging material, processed wood material, and bamboo and rattan products’ (FAO 2021).

