## Supplemental table S1 for "Cosmopolitan Scolytinae: strong common drivers but too many singularities for accurate prediction"

**Table S-1a.** Main characteristics of the 163 Scolytinae with an invasion history. Biological traits

| Species | Species code | Main references on the species[1] | Tribe | Feeding[2] habits | Inbreeding (Y/N) | Polyphagy[3] | Aggregation pheromones[4] | Primary attractants[5] | conifer / non-conifer / both[6] |
|---|---|---|---|---|---|---|---|---|---|
| *Pagiocerus frontalis* (Fabricius) | PAGIFR | WB; Bright & Peck 1998 | Bothrosternini | S | N | 4 | 0 | 0 | 2 |
| *Gnathotrichus materiarius* (Fitch) | GNAHMA | ATK; Kirkendall and Faccoli 2010; Smith & Hulcr (2015); Inward (2020) | Corthylini | A | N | 1 | 2 | 2 | 1 |
| *Monarthrum mali* (Fitch) | MNTHMA | ATK; Kirkendall and Faccoli 2010; Smith & Hulcr (2015) | Corthylini | A | N | 12 | 0 | 2 | 2 |
| *Pityophthorus juglandis* Blackman | PITOJU | ATK; Montecchio et Faccoli 2014; Smith & Hulcr (2015) | Corthylini | B | N | 1 | 2 | 2 | 2 |
| *Pityophthorus solus* (Blackman) | PITOSO | ATK; WB; Goldarazena et al. 2014 | Corthylini | B | N | 1 | 1 | 1 | 1 |
| *Cryphalus pallidus* Eichhoff | cryphpa | CABI Invasive Species Compendium; Johnson et al 2017; Beaver 1987b | Cryphalini | B | N | 2 | 1 | 1 | 2 |
| *Cryphalus wapleri* Eichhoff | CRYHWA | WB; Brockerhoff et al. 2006 | Cryphalini | B | N | 2 | 1 | 1 | 2 |
| *Cryptocarenus heveae* (Hagedorn) | cryphev | ATK; | Corthylini | B | N | 17 | 0 | 2 | 2 |
| *Cryphalus scabricollis* (Eichhoff) | CRYHSC | WB; Kirkendall and Faccoli 2010; Faccoli et al. 2016; Gaaliche et al., 2018; Mifsud & Knizek 2009. | Cryphalini | B | N | 7 | 1 | 1 | 2 |
| *Cryphalus dilutus* (Eichhoff) | hypdil | ATK | Cryphalini | B | N | 2 | 0 | 2 | 2 |
| *Cryphalus mangiferae* (Stebbing) | HYPCMA | ATK; Haack 2001; Beaver 1987b | Cryphalini | B | N | 1 | 0 | 2 | 2 |
| *Hypothenemus africanus* (Hopkins) | HYOTAF | ATK; Haack 2001; Jordal & Kirkendall 1998 | Trypophloeini | H | Y | 4 | 0 | 1 | 2 |
| *Hypothenemus areccae* (Hornung) | HYOTSU | ATK; Haack 2001; padil 2019; Nakagawa et al. 2003 | Trypophloeini | S | Y | 8 | 0 | 2 | 2 |
| *Hypothenemus birmanus* (Eichhoff) | HYOTBI | ATK; Haack 2001; Bright & Peck 1998 | Trypophloeini | B | Y | 29 | 0 | 2 | 2 |
| *Hypothenemus brunneus* Hopkins | HYOTBR | ATK; Haack 2001; Bright & Peck 1998 | Trypophloeini | B | Y | 30 | 0 | 2 | 2 |
| *Hypothenemus californicus* Hopkins | HYOTCA | ATK; Haack 2001 | Trypophloeini | B | Y | 23 | 0 | 2 | 2 |
| *Hypothenemus columbi* Hopkins | HYOTCO | ATK; Haack 2001 | Trypophloeini | B | Y | 22 | 0 | 2 | 2 |

[1] ATK: Atkinson 2022; WB: Wood and Bright. 1992. Note that Bright (2021) was not used for this study because the data search ended in December 2020.

[2] Feeding habits: A: ambrosia beetles (xylomycetophagous); B: bark beetles (phloeophagous); HER: herbiphagous; SPM: spermatophagous

[3] Number of host-plant families (from ATK and WB).

[4] Source for pheromones: El-Sayed (2018). 2 = pheromone(s) identified in the species; 1 = pheromone(s) in at least one other species in the genus; 0 = no pheromone identified or unknown. NB: only semiochemicals identified as pheromones in El-Sayed (2018) are considered; attractants, allomones and synomones are not. For *Euwallacea* spp., the reference is Cooperband et al. (2017)

[5] Sources for primary attractants: main references on the species. 2 = attractant(s) identified in the species; 1 = attractant(s) for at least one other species in the genus; 0 = no attractant(s) identified, or unknown.

[6] 1 = conifers ; 2 = non-conifers ; 3 = non-conifers and conifers.

**Table S-1a.** Main characteristics of the 163 Scolytinae with an invasion history. Biological traits

| Species | Species code | Main references on the species[1] | Tribe | Feeding[2] habits | Inbreeding (Y/N) | Polyphagy[3] | Aggregation pheromones[4] | Primary attractants[5] | conifer / non-conifer / both[6] |
|---|---|---|---|---|---|---|---|---|---|
| *Hypothenemus crudiae* (Panzer) | HYOTHI | ATK; Haack 2001; Jordal & Kirkendall 1998 | Trypophloeini | H | Y | 57 | 0 | 2 | 3 |
| *Hypothenemus elephas* (Eichhoff) | hypele | ATK | Trypophloeini | B | Y | 1 | 0 | 2 | 2 |
| *Hypothenemus erectus* LeConte | HYOTER | ATK; Haack 2001 | Trypophloeini | B | Y | 32 | 0 | 2 | 2 |
| *Hypothenemus eruditus* Westwood | HYOTEU | ATK; Mifsud & Knizek 2009; Kirkendall and Faccoli 2010; Jordal & Kirkendall 1998 | Trypophloeini | H | Y | 65 | 0 | 2 | 3 |
| *Hypothenemus hampei* (Ferrari) | STEHHA | ATK; Jaramillo et al. 2013 | Trypophloeini | S | Y | 1 | 0 | 2 | 2 |
| *Hypothenemus javanus* (Eggers) | HYOTJA | ATK; Haack 2001; Beaver 1987b | Trypophloeini | B | Y | 19 | 0 | 2 | 2 |
| *Hypothenemus leprieuri* ( Perris) | HYOTLE | WB; Mifsud & Knizek 2009 | Trypophloeini | B | Y | 1 | 0 | 1 | 2 |
| *Hypothenemus obscurus* (Fabricius) | HYOTOB | ATK; Haack 2001 | Trypophloeini | B | Y | 11 | 0 | 1 | 2 |
| *Hypothenemus plumeriae* (Nordlinger) | STEHPL | ATK | Trypophloeini | B | Y | 14 | 0 | 2 | 2 |
| *Hypothenemus pubescens* Hopkins | HYOTPB | ATK | Trypophloeini | H | Y | 2 | 0 | 2 | 2 |
| *Hypothenemus seriatus* (Eichhoff) | STEHSE | ATK; Jordal & Kirkendall 1998 | Trypophloeini | H | Y | 57 | 0 | 2 | 3 |
| *Hypothenemus setosus* (Eichhoff) | HYOTSE | ATK; Haack 2001; Jordal & Kirkendall 1998 | Trypophloeini | H | Y | 9 | 0 | 2 | 2 |
| *Eidophelus jalapae (Letzner)* | scojal | ATK | Ernoporini | B | N | 9 | 0 | 2 | 2 |
| *Aphanarthrum* (*Coleobothrus*) *alluaudi* Peyerimhoff | aphall | WB; Kirkendall et al. 2015 | Crypturgini | H | N | 1 | 0 | 0 | 2 |
| *Aphanarthrum affine* Wollaston | aphaff | WB; Israelson 1972 | Crypturgini | H | N | 1 | 0 | 0 | 2 |
| *Aphanarthrum bicolor* Wollaston | aphbic | WB; Israelson 1972 | Crypturgini | H | N | 1 | 0 | 0 | 2 |
| *Aphanarthrum mairei* Peyerimhoff | aphmai | WB; Israelson 1972 | Crypturgini | H | N | 1 | 0 | 0 | 2 |
| *Aphanarthrum piscatorium* Wollaston | aphpis | WB; Israelson 1972 | Crypturgini | H | N | 1 | 0 | 0 | 2 |
| *Crypturgus cylindricollis* Eggers | CRYUCY | Mifsud & Knizek 2009 | Crypturgini | B | N | 1 | 0 | 1 | 1 |
| *Crypturgus numidicus* Ferrari | CRYUNU | WB; Mifsud & Knizek 2009 | Crypturgini | B | N | 1 | 0 | 1 | 1 |
| *Crypturgus pusillus* (Gyllenhal) | CRYUPU | ATK; Haack 2001 | Crypturgini | B | N | 1 | 0 | 2 | 1 |
| *Coccotrypes aciculatus* Schedl | cocoac | ATK | Dryocoetini | S | Y | 1 | 0 | 1 | 2 |
| *Coccotrypes advena* Blandford | COCOAD | WB; Haack 2001; Brockerhoff et al. 2006; Nakagawa et al. 2003; Jordal & Kirkendall 1998 | Dryocoetini | S | Y | 20 | 0 | 1 | 3 |
| *Coccotrypes carpophagus* (Hornung) | COCOCA | ATK; Haack 2001; Bright & Peck 1998 | Dryocoetini | S | Y | 12 | 0 | 2 | 2 |
| *Coccotrypes cyperi* (Beeson) | COCOCY | ATK; Haack 2001; Jordal & Kirkendall 1998 | Dryocoetini | H | Y | 24 | 0 | 2 | 2 |
| *Coccotrypes dactyliperda* Fabricius | COCODA | WB; Haack 2001; Kirkendall et Faccoli 2010; Brockerhoff et al. 2006; Kirkendall 2018; Mifsud & Knizek 2009; Beaver 1987b | Dryocoetini | S | Y | 6 | 0 | 2 | 2 |

**Table S-1a.** Main characteristics of the 163 Scolytinae with an invasion history. Biological traits

| Species | Species code | Main references on the species[1] | Tribe | Feeding[2] habits | Inbreeding (Y/N) | Polyphagy[3] | Aggregation pheromones[4] | Primary attractants[5] | conifer / non-conifer / both[6] |
|---|---|---|---|---|---|---|---|---|---|
| *Coccotrypes distinctus* (Motschulsky) | COCODI | ATK; Haack 2001; Brockerhoff et al. 2006; Kirkendall 2018 | Dryocoetini | S | Y | 4 | 0 | 2 | 2 |
| *Coccotrypes rhizophorae* (Hopkins) | COCORH | ATK; Wood 1982; Haack 2001 | Dryocoetini | S | Y | 5 | 0 | 1 | 2 |
| *Coccotrypes robustus* Eichhoff | COCORO | ATK; Wood 1982; Haack 2001 | Dryocoetini | S | Y | 1 | 0 | 1 | 2 |
| *Coccotrypes rutschuruensis* Eggers | COCORU | ATK; Wood 2007; Haack 2001 | Dryocoetini | S | Y | 3 | 0 | 1 | 2 |
| *Coccotrypes vulgaris* (Eggers) | COCOVU | ATK; Haack 2001 | Dryocoetini | B | Y | 8 | 0 | 1 | 2 |
| *Cyrtogenius luteus* (Blanford) | CYRGLU | WB; Fletchmann and Atkinson 2018; Faccoli et al. 2012; Gomez et al. 2012; Kirkendall et al. 2015 | Dryocoetini | B | N | 1 | 0 | 2 | 1 |
| *Dactylotrypes longicollis* (Wollaston) | DACPLO | ATK; ; Kirkendall et al. 2015; Kirkendall and Faccoli 2010; Kirkendall 2018 | Dryocoetini | S | N | 2 | 0 | 2 | 2 |
| *Dryocoetes himalayensis* Strohmeyer | DRYOHI | WB; Kirkendall and Faccoli 2010 | Dryocoetini | B | N | 2 | 1 | 1 | 2 |
| *Thamnurgus characiae* Rosenhauer | THMNCH | WB; Mifsud & Knizek 2009 | Dryocoetini | B | N | 1 | 0 | 0 | 2 |
| *Microborus boops* Blandford | micboop | ATK | Hexacolini | B | N | 1 | 0 | 0 | 2 |
| *Hylastes angustatus* (Herbst) | HYASAN | WB; Schedl 1957, 1965; Tribe 1992 | Hylastini | B | N | 1 | 0 | 1 | 1 |
| *Hylastes ater* (Paykull) | HYASAR | WB; Brockerhoff et al. 2003; Kirkendall 2018 | Hylastini | B | N | 2 | 0 | 2 | 1 |
| *Hylastes linearis* Erichson | HYASLI | WB; Kirkendall 2018 | Hylastini | B | N | 1 | 0 | 1 | 1 |
| *Hylastes opacus* Erichson | HYASOP | ATK; Haack 2001; 2006 | Hylastini | B | N | 1 | 0 | 2 | 1 |
| *Hylastinus obscurus* (Marsham) | HYATOB | ATK; Haack 2001 | Hylastini | B | N | 1 | 0 | 2 | 2 |
| *Hylurgops palliatus* (Gyllenhal) | HYLUPA | ATK; Haack 2001; 2006 | Hylastini | B | N | 1 | 0 | 2 | 1 |
| *Hylesinus toranio* (= *Bostrychus oleiperda*) (Danthione) | HYESOL | ATK; Kirkendall 2018 | Hylesinini | B | N | 1 | 0 | 1 | 2 |
| *Kissophagus hederae* (Schmidt) | KISSHE | WB; Mifsud & Knizek 2009 | Hylesinini | B | N | 1 | 0 | 0 | 2 |
| *Dendroctonus micans* (Kugelann) | DENCMI | WB; Bevan & King 1983 | Hylurgini | B | Y | 1 | 0 | 0 | 1 |
| *Dendroctonus valens* LeConte | DENCVA | ATK; Yan et al. 2005 | Hylurgini | B | N | 1 | 2 | 2 | 1 |
| *Hylurgus ligniperda* (Fabricius) | HYLGLI | ATK; Haack 2001; 2006; Brockerhoff et al. 2006; Kirkendall 2018 | Hylurgini | B | N | 1 | 0 | 2 | 1 |
| *Hylurgus micklitzi* Wachtl | HYLGNI | WB; Mifsud & Knizek 2009 | Hylurgini | B | N | 1 | 0 | 1 | 1 |
| *Pseudohylesinus sericeus* (Mannerheim) | PSDHSE | ATK | Hylurgini | B | N | 1 | 0 | 2 | 1 |
| *Tomicus piniperda* (Linnaeus) | BLASPI | ATK; Haack 2001 | Hylurgini | B | N | 1 | 0 | 2 | 1 |
| *Hypoborus ficus* Erichson | HYPBFI | WB; Israelson 1990 | Hypoborini | B | N | 1 | 0 | 1 | 2 |
| *Liparthrum artemisiae* Wollaston | lipart | WB; Israelson 1990 | Hypoborini | B | N | 1 | 0 | 0 | 2 |
| *Liparthrum bituberculatum* Wollaston | lipbit | WB; Israelson 1990 | Hypoborini | B | N | 1 | 0 | 0 | 2 |
| *Liparthrum curtum* Wollaston | lipcur | WB; Israelson 1990 | Hypoborini | B | N | 3 | 0 | 0 | 2 |
| *Liparthrum inarmatum* Wollaston | lipina | WB; Israelson 1990 | Hypoborini | B | N | 1 | 0 | 0 | 2 |
| *Liparthrum mandibulare* Wollaston | lipman | WB; Israelson 1990 | Hypoborini | B | N | 1 | 0 | 0 | 2 |

**Table S-1a.** Main characteristics of the 163 Scolytinae with an invasion history. Biological traits

| Species | Species code | Main references on the species[1] | Tribe | Feeding[2] habits | Inbreeding (Y/N) | Polyphagy[3] | Aggregation pheromones[4] | Primary attractants[5] | conifer / non-conifer / both[6] |
|---|---|---|---|---|---|---|---|---|---|
| *Liparthrum mori* (Aube) | LPRTMO | WB; Mifsud & Knizek 2009 | Hypoborini | B | N | 2 | 0 | 0 | 2 |
| *Ips calligraphus* (Germar) | IPSXCA | ATK | Ipini | B | N | 1 | 2 | 2 | 1 |
| *Ips cembrae* (Herr) | IPSXCE | WB; Crooke & Bevan 1957 | Ipini | B | N | 1 | 2 | 0 | 1 |
| *Ips grandicollis* (Eichhoff) | IPSXGR | ATK | Ipini | B | N | 1 | 2 | 2 | 1 |
| *Orthotomicus angulatus* (Eichhoff) | ortang | WB | Ipini | B | N | 2 | 1 | 1 | 1 |
| *Orthotomicus caelatus* (Eichhoff) | ORTCCA | ATK; | Ipini | B | N | 1 | 0 | 2 | 1 |
| *Orthotomicus erosus* (Wollaston) | IPSXER | ATK; Haack 2006; Mifsud & Knizek 2009; Tribe 1992 | Ipini | B | N | 1 | 2 | 2 | 1 |
| *Orthotomicus laricis* (Fabricius) | IPSXLC | WB; Kirkendall 2018 | Ipini | B | N | 1 | 0 | 1 | 1 |
| *Orthotomicus proximus* (Eichhoff) | IPSXPR | WB | Ipini | B | N | 1 | 1 | 1 | 1 |
| *Pityogenes bidentatus* (Herbst) | PITYBD | ATK; Haack 2001 | Ipini | B | N | 1 | 2 | 2 | 1 |
| *Pityogenes calcaratus* (Eichhoff) | PITYCC | WB; Mifsud & Knizek 2009 | Ipini | B | N | 1 | 2 | 1 | 1 |
| *Pityogenes chalcographus* (Linnaeus) | PITYCH | WB; Haack 2001 | Ipini | B | N | 1 | 2 | 1 | 1 |
| *Pityokteines curvidens* (Germar) | PITKCU | WB | Ipini | B | N | 1 | 2 | 0 | 1 |
| *Premnobius ambitiosus* (Schaufuss) | preamb | ATK | Ipini | B | Y | 5 | 0 | 1 | 2 |
| *Premnobius cavipennis* Eichhoff | PREBCA | ATK; Haack 2001 | Ipini | B | Y | 32 | 0 | 2 | 2 |
| *Phloeosinus armatus* Reitter | PHLSAR | ATK; Haack 2001 | Phloeosinini | B | N | 1 | 0 | 1 | 1 |
| *Phloeosinus cupressi* Hopkins | PHLSCU | ATK; Neumann 1987; Brockerhoff et al. 2006; Wood 1977? | Phloeosinini | B | N | 2 | 0 | 2 | 1 |
| *Phloeosinus rudis* Blandford | PHLSRD | WB; Kirkendall and Faccoli 2010 | Phloeosinini | B | N | 1 | 0 | 1 | 1 |
| *Phloeosinus thujae* (Perris) | PHLSTH | WB; Mifsud & Knizek 2009 | Phloeosinini | B | N | 1 | 0 | 1 | 1 |
| *Phloeotribus liminaris* (Harris) | PHLBLI | ATK; Kirkendall and Faccoli 2010 | Phloeotribini | B | N | 2 | 0 | 2 | 2 |
| *Phloeotribus scarabaeoides* (Bernard) | PHLBOL | WB | Phloeotribini | B | N | 2 | 0 | 2 | 2 |
| *Polygraphus poligraphus* (Linnaeus) | POLGPO | WB; EPPO GD | Polygraphini | B | N | 1 | 2 | 1 | 1 |
| *Polygraphus proximus* Blandford | POLGPR | WB; EPPO GD | Polygraphini | B | N | 1 | 2 | 1 | 1 |
| *Polygraphus rufipennis* (Kirby) | POLGRU | WB; EPPO GD | Polygraphini | B | N | 1 | 2 | 1 | 1 |
| *Scolytus amygdali* Guerin-Meneville | SCOLAM | WB; Mifsud & Knizek 2009 | Scolytini | B | N | 1 | 2 | 1 | 2 |
| *Scolytus dimidiatus* Chapuis | scodim | ATK | Scolytini | B | N | 2 | 1 | 1 | 2 |
| *Scolytus kirschi* Skahtzky | SCOLKI | WB; Six et al. 2005 | Scolytini | B | N | 4 | 1 | 1 | 2 |
| *Scolytus mali* (Bechstein) | SCOLMA | ATK; Haack 2001 | Scolytini | B | N | 2 | 1 | 2 | 2 |
| *Scolytus multistriatus* (Marsham) | SCOLMU | ATK; Haack 2001; Brockerhoff et al. 2006; Fauna Europaea 2019; Kirkendall 2018 | Scolytini | B | N | 3 | 2 | 2 | 2 |
| *Scolytus rugulosus* (Muller) | SCOLRU | ATK; Haack 2001; Mifsud & Knizek 2009; Kirkendall 2018 | Scolytini | B | N | 1 | 1 | 1 | 2 |
| *Scolytus schevyrewi* Semenov | SCOLSH | ATK; Haack 2001; Lee et al., 2009 | Scolytini | B | N | 5 | 1 | 2 | 2 |
| *Scolytus sulcifrons* Rey | SCOLSU | Mifsud & Knizek 2009 | Scolytini | B | N | 2 | 1 | 1 | 2 |

**Table S-1a.** Main characteristics of the 163 Scolytinae with an invasion history. Biological traits

| Species | Species code | Main references on the species[1] | Tribe | Feeding[2] habits | Inbreeding (Y/N) | Polyphagy[3] | Aggregation pheromones[4] | Primary attractants[5] | conifer / non-conifer / both[6] |
|---|---|---|---|---|---|---|---|---|---|
| *Scolytoplatypus tycon* Blandford | scotyc | WB; Beaver and Gebhardt 2006; Shapovalov et al. 2010 | Scolytoplatypodini | A | N | 11 | 0 | 0 | 3 |
| *Amasa truncatus* (*Xyleborus truncatus*) (Erichson) | XYLBTR | WB; Brockerhoff et al. 2006; Kirkendall 2018; Gomez et al. 2017 | Xyleborini | A | Y | 2 | 0 | 2 | 3 |
| *Ambrosiodmus compressus* (Lea) | AMBDCO | WB; Brockerhoff et al. 2006 | Xyleborini | A | Y | 6 | 0 | 2 | 3 |
| *Ambrosiodmus lewisi* (Blandford) | AMBDLE | WB; Haack 2001; 2006 | Xyleborini | A | Y | 15 | 0 | 2 | 2 |
| *Ambrosiodmus minor* (Stebbing) | AMBDMI | ATK | Xyleborini | A | Y | 5 | 0 | 1 | 2 |
| *Ambrosiodmus obliquus* (LeConte) | AMBDOB | ATK; Faccoli et al. 2009 | Xyleborini | A | Y | 17 | 0 | 2 | 2 |
| *Ambrosiodmus rubricollis* (Eichhoff) | AMBDRU | HB; Haack 2001; Kirkendall et Faccoli 2010 | Xyleborini | A | Y | 17 | 0 | 2 | 2 |
| *Ambrosiophilus* (*Xyleborus*) *atratus* (Eichhoff) | XYLBAT | ATK; Kirkendall and Faccoli 2010; Haack 2001; 2006 | Xyleborini | A | Y | 4 | 0 | 2 | 3 |
| *Ambrosiophilus nodulosus* (Eggers) | AMBHNO | ATK; Smith and Cognato 2015 ; Smith et al. 2017 | Xyleborini | A | Y | 3 | 0 | 1 | 2 |
| *Anisandrus* (*Xyleborus*) *dispar* (Fabricius) | XYLBDI | ATK; Haack 2001 | Xyleborini | A | Y | 17 | 0 | 2 | 3 |
| *Anisandrus maiche* (Kurentzov) | ANIDMA | ATK; Rabaglia et al. 2009; EPPO 2013 | Xyleborini | A | Y | 5 | 0 | 2 | 2 |
| *Cnestus* (*Xylosandrus*) *mutilatus* (Blandford | XYLSMU | ATK; Haack 2001; 2006 | Xyleborini | A | Y | 13 | 0 | 2 | 3 |
| *Cyclorhipidion bodoanum* (*Xyleborus californicus*) (Reitter) | XYLBCA | ATK; Haack 2001; Kirkendall and Faccoli 2010; Inward (in press) | Xyleborini | A | Y | 3 | 0 | 2 | 2 |
| *Cyclorhipidion fukiense* (Eggers) | CYCRFU | ATK; Hoebeke et al. 2018 | Xyleborini | A | Y | 1 | 1 | 2 | 2 |
| *Cyclorhipidion pelliculosum* (Eichhoff) | XYLBPL | ATK; Atkinson et al. 1990 | Xyleborini | A | Y | 3 | 0 | 2 | 2 |
| *Dryocoetoides cristatus* (Fabricius) | drycris | WB; Zanuncio et al 2005; | Xyleborini | A | Y | 4 | 0 | 1 | 2 |
| *Dryoxylon onoharaense* (Murayama) | DRYXON | ATK; Haack 2001 | Xyleborini | A | Y | 4 | 0 | 0 | 2 |
| *Eccoptopterus spinosus* (Olivier) | ECCOSI | WB; Beaver 1988 | Xyleborini | A | Y | 10 | 0 | 2 | 2 |
| *Euwallacea* (*Xyleborus*) *piceus* (Motschulsky) | EUWAPI | WB; Beaver 1988 | Xyleborini | A | Y | 26 | 1 | 1 | 3 |
| *Euwallacea (Xyleborus) similis (Ferrari)* | XYLBSI | Atkinson 2021; Haack 2001; 2006; O'Donnell et al. 2016; Beaver & Liu 2016 | Xyleborini | A | Y | 5 | 1 | 2 | 3 |
| *Euwallacea fornicatus* (Eichhoff) | EUWAWH | ATK; Haack 2006; Mendel et al., 2012; Paap et al. 2018; Beaver & Liu 2016. | Xyleborini | A | Y | 18 | 2 | 2 | 2 |
| *Euwallacea interjectus* (Blandford) | XYLBIN | ATK; Cognato et al. 2015 | Xyleborini | A | Y | 19 | 1 | 2 | 3 |
| *Euwallacea kuroshio* Gomez and Hulcr | EUWAKU | ATK; Gomez et al 2018b | Xyleborini | A | Y | 1 | 1 | 1 | 2 |
| *Euwallacea perbrevis (Schedl, 1951)* | EUWAPE | ATK; Smith et al 2019 | Xyleborini | A | Y | 17 | 2 | 2 | 2 |
| *Euwallacea validus* (Eichhoff) | XYLBVA | ATK; Haack 2001 | Xyleborini | A | Y | 18 | 1 | 2 | 3 |
| *Microperus* (*Coptodryas*) *eucalypticus* (Schedl) | MIPREU | Morgan 1967; Brockerhoff et al. 2006; Beaver & Liu 2016 | Xyleborini | A | Y | 6 | 0 | 0 | 2 |

**Table S-1a.** Main characteristics of the 163 Scolytinae with an invasion history. Biological traits

| Species | Species code | Main references on the species[1] | Tribe | Feeding[2] habits | Inbreeding (Y/N) | Polyphagy[3] | Aggregation pheromones[4] | Primary attractants[5] | conifer / non-conifer / both[6] |
|---|---|---|---|---|---|---|---|---|---|
| *Microperus quercicola* (Eggers) | micquer | Mandelshtam et al 2018; Lantschner et al 2020 | Xyleborini | A | Y | 4 | 0 | 0 | 2 |
| *Planiculus (Euwallacea) bicolor* (Blandford) | euwabic | Beaver 1988 | Xyleborini | A | Y | 12 | 1 | 1 | 2 |
| *Theoborus ricini* (Eggers) | theori | ATK | Xyleborini | A | Y | 13 | 0 | 2 | 2 |
| *Truncaudum* (*Cyclorhipidion*) *agnatum* (Eggers) | TRUCAG | WB; Beaver & Liu 2016 | Xyleborini | A | Y | 4 | 0 | 2 | 2 |
| *Xyleborinus andrewesi* (Blandford) | XYBIAN | ATK; Okins & Thomas 2010; Beaver 1988 | Xyleborini | A | Y | 14 | 0 | 2 | 2 |
| *Xyleborinus artestriatus* (Eichhoff) | XYBIAR | ATK; Cognato et al. 2013 | Xyleborini | A | Y | 7 | 0 | 2 | 2 |
| *Xyleborinus attenuatus* (Blandford) | XYBIAL | ATK; Haack 2001; 2006; Kirkendall et Faccoli 2010 | Xyleborini | A | Y | 7 | 0 | 2 | 2 |
| *Xyleborinus exiguus* (Walker) | XYBIEX | ATK | Xyleborini | A | Y | 13 | 0 | 1 | 2 |
| *Xyleborinus gracilis* Eichhoff | XYBIGR | ATK; Bright & Peck 1998 | Xyleborini | A | Y | 14 | 0 | 2 | 2 |
| *Xyleborinus octiesdentatus* (Murayama) | XYBIOC | ATK; Rabaglia et al. 2010 | Xyleborini | A | Y | 3 | 0 | 2 | 2 |
| *Xyleborinus saxeseni* (Ratzeburg) | XYLBSA | ATK; Haack 2001; Brockerhoff et al. 2006; Mifsud & Knizek 2009; Beaver & Liu 2016 | Xyleborini | A | Y | 24 | 0 | 2 | 3 |
| *Xyleborus affinis* Eichhoff | XYLBAF | ATK; Kirkendall & Faccoli 2010; Beaver & Liu 2016; Beaver 1988; Beaver & Liu 2016 | Xyleborini | A | Y | 47 | 0 | 2 | 3 |
| *Xyleborus africanus* Eggers | xylafr | WB; Beaver 1988 | Xyleborini | A | Y | 7 | 0 | 1 | 2 |
| *Xyleborus atratus* Eichhoff | xylatr | WB; Faccoli 2008 | Xyleborini | A | Y | 15 | 0 | 2 | 3 |
| *Xyleborus bispinatus* Eichhoff | XYLBBI | ATK; Faccoli et al. 2016 | Xyleborini | A | Y | 6 | 0 | 2 | 2 |
| *Xyleborus ferrugineus* (Fabricius) | XYLBFE | ATK; Beaver & Liu 2016; Bright & Peck 1998; Beaver 1988; Beaver & Liu 2016 | Xyleborini | A | Y | 33 | 0 | 2 | 3 |
| *Xyleborus glabratus* Eichhoff | XYLBGR | ATK; Haack 2001; 2006 | Xyleborini | A | Y | 4 | 0 | 2 | 3 |
| *Xyleborus monographus* (Fabricius) | XYLBMO | ATK; Rabaglia et al. 2020; Smith et al 2020 | Xyleborini | A | Y | 6 | 0 | 2 | 2 |
| *Xyleborus perforans* (Wollaston) | XYLBPE | WB; Beaver & Liu 2016; Beaver 1988 | Xyleborini | A | Y | 28 | 0 | 2 | 2 |
| *Xyleborus pfeilii* (Ratzeburg) | XYLBPF | ATK; Haack 2001, 2006; Kirkendall et Faccoli 2010 | Xyleborini | A | Y | 13 | 0 | 2 | 3 |
| *Xyleborus seriatus* Blandford | XYLBSE | ATK; Haack 2006 | Xyleborini | A | Y | 12 | 0 | 2 | 3 |
| *Xyleborus spinulosus* Blandford | XYLBSN | ATK; Bright & Peck 1998 | Xyleborini | A | Y | 13 | 0 | 2 | 3 |
| *Xyleborus volvulus* (Fabricius) | xylvol | ATK; Beaver 1988; Bright & Peck 1998 | Xyleborini | A | Y | 25 | 0 | 2 | 3 |
| *Xylosandrus* (*Apoxyleborus*) *mancus* (Blandford) | xylman | WB; Beaver 1988 | Xyleborini | A | Y | 14 | 0 | 2 | 2 |
| *Xylosandrus amputatus* (Blandford) | XYLSAM | ATK; Cognato et al. 2011 | Xyleborini | A | Y | 4 | 0 | 2 | 2 |

**Table S-1a.** Main characteristics of the 163 Scolytinae with an invasion history. Biological traits

| **Species** | **Species code** | **Main references on the species**[1] | **Tribe** | **Feeding**[2] **habits** | **Inbreeding (Y/N)** | **Polyphagy**[3] | **Aggregation pheromones**[4] | **Primary attractants**[5] | **conifer / non-conifer / both**[6] |
|---|---|---|---|---|---|---|---|---|---|
| *Xylosandrus compactus* (Eichhoff) | XYLSCO | ATK; Haack 2001; Garonna et al. 2012; Beaver 1988; Beaver & Liu 2016 | Xyleborini | A | Y | 28 | 0 | 2 | 2 |
| *Xylosandrus crassiusculus* (Motschulsky) | XYLSCR | ATK; Haack 2001; Kirkendall and Faccoli 2010; Kirkendall 2018; padil 2019; Beaver 1988; Beaver & Liu 2016; EPPO 2019 | Xyleborini | A | Y | 51 | 0 | 2 | 3 |
| *Xylosandrus germanus* (Blandford) | XYLSGE | ATK; Haack 2001; Kirkendall et Faccoli 2010; | Xyleborini | A | Y | 29 | 0 | 2 | 3 |
| *Xylosandrus morigerus* (Blandford) | XYLSMO | ATK; Kirkendall et Faccoli 2010; Beaver 1988; Beaver & Liu 2016 | Xyleborini | A | Y | 39 | 0 | 2 | 2 |
| *Xylosandrus pseudosolidus* (Schedl) | XYLSPS | WB; Brockerhoff et al. 2006 | Xyleborini | A | Y | 4 | 0 | 1 | 3 |
| *Xyloterinus politus* (Say) | xypol | WB; Dodelin & Saurat 2017 | Xyleborini | A | Y | 7 | 0 | 2 | 2 |
| *Trypodendron domesticum* (Linnaeus) | TRYDDO | ATK; Haack 2001; Fauna Europaea 2019 | Xyloterini | A | N | 8 | 2 | 2 | 2 |

**Table S-1b.** Main characteristics of the 163 Scolytinae with an invasion history. Impact and landmasses colonised

| Species | Spec code | Impact on plant health[7] | References on impact | Comments on impact | Nb of land masses | Landmasses[8] |
|---|---|---|---|---|---|---|
| *Amasa truncatus* (*Xyleborus truncatus*) (Erichson) | XYLBTR | 1 | damage | Borer of dead exotics and natives, also in live eucalypts | 4 | AUS; TAS; SAM; NZL |
| *Ambrosiodmus compressus* (Lea) | AMBDCO | 1 | Brockerhoff & Bain (2000) | Borer of dead exotics and natives, also in live eucalypts | 2 | AUS; NZL |
| *Ambrosiodmus lewisi* (Blandford) | AMBDLE | 0 | Haack 2006 | From dead oak branches | 9 | ASI; JPN; LKA; TWN; BOR; JAW; SUM; PHL; NAM |
| *Ambrosiodmus minor* (Stebbing) | AMBDMI | 0 | Hulcr et al. 2018 | Wood decay | 2 | NAM; ASI |
| *Ambrosiodmus obliquus* (LeConte) | AMBDOB | 0 | Faccoli et al. 2009 | | 7 | AFR; CAM; HIS; GLP; PRI; NAM; SAM |
| *Ambrosiodmus rubricollis* (Eichhoff) | AMBDRU | 1 | Kirkendall & Faccoli (2010) | On a live *Aesculus hippocastaneum* | 7 | NAM; EUR; ASI; BON; JPN; TWN; AUS |
| *Ambrosiophilus* (*Xyleborus*) *atratus* (Eichhoff) | XYLBAT | 0 | Rabaglia 2006 | | 10 | ASI; JAW; SUM; PHL; JPN; NAM; NWG; EUR; TWN; IDN |
| *Ambrosiophilus nodulosus* (Eggers) | AMBHNO | 0 | | No data on damage to plant health | 2 | ASI; NAM |
| *Anisandrus* (*Xyleborus*) *dispar* (Fabricius) | XYLBDI | 1 | CABI 2021b | The beetle preferentially attacks stressed trees. | 4 | ASI; NAM; EUR; GBR |
| *Anisandrus maiche* (Kurentzov) | ANIDMA | 0 | Terekhova & Skrylnik (2012) | Drying out trees and fresh deadwood trees | 3 | ASI; NAM ; EUR |
| *Aphanarthrum* (*Coleobothrus*) *alluaudi* Peyerimhoff | aphall | 0 | | Fungi: Kolarik et al (2007). Impact: No data on damage to plant health | 2 | AFR; CNY |
| *Aphanarthrum affine* Wollaston | aphaff | 0 | | Fungi: Kolarik et al (2007). Impact: No data on damage to plant health | 3 | AFR; EUR; CNY |
| *Aphanarthrum bicolor* Wollaston | aphbic | 0 | | Fungi: Kolarik et al (2007). Impact: No data on damage to plant health | 2 | CNY; MDR |
| *Aphanarthrum mairei* Peyerimhoff | aphmai | 0 | | Fungi: Kolarik et al (2007). Impact: No data on damage to plant health | 3 | AFR; CNY; MDR |
| *Aphanarthrum piscatorium* Wollaston | aphpis | 0 | | Fungi: Kolarik et al (2007). Impact: No data on damage to plant health | 2 | CNY; MDR |
| *Cnestus* (*Xylosandrus*) *mutilatus* (Blandford | XYLSMU | 0 | CABI 2018 | https://www.cabi.org/isc/datasheet/57239 | 8 | ASI; NAM; LKA; JPN; OKI; TWN; IDN; NWG |
| *Coccotrypes aciculatus* Schedl | cocoac | 0 | | No data on damage to plant health | 6 | CAM; BRB; CUB; NAM; NWG; SAM |

[7] 0 = no documented impact; 1 = indications of impact, with some uncertainties; 2 = known substantial impact

[8] See the acronyms in Table S-2. Landmasses: all areas isolated by a sea or an ocean (islands and continents).

**Table S-1b.** Main characteristics of the 163 Scolytinae with an invasion history. Impact and landmasses colonised

| Species | Spec code | Impact on plant health[7] | References on impact | Comments on impact | Nb of land masses | Landmasses[8] |
|---|---|---|---|---|---|---|
| *Coccotrypes advena* Blandford | COCOAD | 0 | Seybold et al. 2016 | *Coccotrypes advena* Blandford is thought to be native to Asia (Indonesia) and breeds in either bark or large seeds of a variety of tropical hosts | 16 | CUB; JPN; LKA; AUS; NZL; FJI; BOR; JAW; SUM; HAW; FSM; NWG; NAM; PHL; WSM; SAM |
| *Coccotrypes carpophagus* (Hornung) | COCOCA | 1 | Rodriguez et al. 2014 | "Janzen [14] found up to 99% of *Euterpe globosa* seeds damaged by *Coccotrypes carpophagus* in Puerto Rico" | 30 | AFR; AZO; CNY; MDG; REU; SYC; ASI; IDN; PHL; LKA; CAM; BMU; CUB; GRD; GLP; JAM; MSR; PRI; HIS; VIL; GBR; EUR; JPN; NAM; AUS; GUM; HAW; SAM; TTO; GAL |
| *Coccotrypes cyperi* (Beeson) | COCOCY | 0 | Kirkendall 2018; Beaver 1987b | "This polyphagous *Coccotrypes* is widely distributed in tropical and subtropical environments around the world, and breeds in everything from seeds and twigs to under bark of branches" | 21 | ASI; IDN; LKA; CAM; GLP; JAM; MTQ; PRI; HIS; NAM; AUS; COK; FJI; FSM; WSM; TAH; TON; HAW; SAM; TTO; SYC |
| *Coccotrypes dactyliperda* Fabricius | COCODA | 2 | Rodriguez et al. 2014 | "Blumberg [33] quoted a 30–40% yield loss of unripe *P. dactylifera* dates attacked by C. dactyliperda in Israel" | 25 | AFR; CNY; MDG; MDR; ASI; IDN; CAM; BHS; CUB; JAM; PRI; HIS; EUR; JPN; NAM; AUS; NWG; NZL; SLB; HAW; GAL; TTO; SAM; MLT; SYC |
| *Coccotrypes distinctus* (Motschulsky) | COCODI | 0 | | No data on damage to plant health | 14 | LKA; CAM; JAM; PRI; HIS; VIL; BMU; NAM; GUM; FSM; NWG; HAW; SAM; NZL |
| *Coccotrypes rhizophorae* (Hopkins) | COCORH | 2 | Sousa et al. 2003 | "Mortality due to beetle attack increased linearly from an average of 10% inside a gap to 72% at 20 m into the forest" | 6 | ASI; JAW; SUM; IDN; NAM; GAL |
| *Coccotrypes robustus* Eichhoff | COCORO | 0 | | No data on damage to plant health | 3 | PRI; CUB; NAM |
| *Coccotrypes rutschuruensis* Eggers | COCORU | 0 | | No data on damage to plant health | 3 | AFR; NAM; SAM |
| *Coccotrypes vulgaris* (Eggers) | COCOVU | 0 | Beaver 1976 | | 11 | ASI; IDN; LUZ; LKA; CAM; PRI; NAM; FJI; NWG; WSM; SAM |
| *Cryphalus pallidus* Eichhoff | cryphpa | 0 | Johnson et al 2017 | in Lantschner et al 2020 | 4 | AFR; EUR; SYC; MDG |
| *Cryphalus wapleri* Eichhoff | CRYHWA | 0 | | No data on damage to plant health | 3 | AUS; NZL; NWG |
| *Cryptocarenus heveae* (Hagedorn) | cryphev | 0 | Wood 1977 | "It infests the pith of small, broken, or unthrifty stems of a wide variety of trees, shrubs, and woody vines." | 10 | AFR; CAM; CUB; GLP; JAM; PRI; HIS; NAM; SAM; TTO |
| *Crypturgus cylindricollis* Eggers | CRYUCY | 0 | | No data on damage to plant health | 3 | EUR; ASI; MLT |
| *Crypturgus numidicus* Ferrari | CRYUNU | 0 | | No data on damage to plant health | 7 | EUR; COR; SAR; SIC; AFR; ASI; MLT |

**Table S-1b.** Main characteristics of the 163 Scolytinae with an invasion history. Impact and landmasses colonised

| Species | Spec code | Impact on plant health[7] | References on impact | Comments on impact | Nb of land masses | Landmasses[8] |
|---|---|---|---|---|---|---|
| *Crypturgus pusillus* (Gyllenhal) | CRYUPU | 0 | | No data on damage to plant health | 7 | AFR; ASI; EUR; GBR; TWN; JPN; NAM |
| *Cyclorhipidion bodoanum* (*Xyleborus californicus*) (Reitter) | XYLBCA | 0 | Inward 2019 | No data on damage to plant health | 4 | ASI; NAM; EUR; GBR |
| *Cyclorhipidion fukiense* (Eggers) | CYCRFU | 0 | | No data on damage to plant health | 3 | ASI; TWN; NAM |
| *Cyclorhipidion pelliculosum* (Eichhoff) | XYLBPL | 0 | | No data on damage to plant health | 3 | ASI; JPN; NAM |
| *Cyrtogenius luteus* (Blanford) | CYRGLU | 0 | | No data on damage to plant health | 14 | ASI; JPN; TWN; JAW; PHL; FSM; BIS; CRL; GIL; MAR; WSM; NWG; SAM; EUR |
| *Dactylotrypes longicollis* (Wollaston) | DACPLO | 0 | | No data on damage to plant health | 5 | MDR; CNY; EUR; NAM; SAM |
| *Dendroctonus micans* (Kugelann) | DENCMI | 2 | Grégoire 1988 | | 4 | ASI; SAK; EUR; GBR |
| *Dendroctonus valens* LeConte | DENCVA | 2 | Liu et al. 2013 | Fungi: Lu et al. 2009 | 3 | CAM; NAM; ASI |
| *Dryocoetes himalayensis* Strohmeyer | DRYOHI | 0 | Foit et al. 2017 | | 2 | ASI; EUR |
| *Dryocoetoides cristatus* (Fabricius) | drycris | 0 | | No data on damage to plant health | 4 | AFR; STP; TTO; SAM |
| *Dryoxylon onoharaense* (Murayama) | DRYXON | 0 | | No data on damage to plant health | 2 | NAM; JPN |
| *Eccoptopterus spinosus* (Olivier) | ECCOSI | 0 | Beaver 1987 | No data on damage to plant health | 19 | AFR; SYC; CNY; FER; ASI; AND; JPN; LKA; TWN; AUS; BOR; CEL; JAW; SUM; MDG; NWG; PHL; REU; TON |
| *Euwallacea* (*Xyleborus*) *piceus* (Motschulsky) | EUWAPI | 1 | CABI 2021d | "It attacks dying, dead and recently felled trees. It has not been found in living trees" | 16 | SYC; AFR; AND; LKA; ASI; TWN; AUS; FJI; BOR; JAW; MEN; MDG; FSM; NWG; PHL; WSM |
| *Euwallacea (Xyleborus) similis (Ferrari)* | XYLBSI | 1 | CABI 2021k | « Attacks by *X. similis* are normally secondary on stressed, dying or dead trees. However, the species could become a pest in reforestation projects or in plantations" | 26 | AFR; MDG; MUS; SYC; CCK; ASI; IDN; PHL; LKA; TWN; JPN; NAM; AUS; CRL; CXR; FJI; GUM; KIR; MHL; FSM; NWG; SLB; TAH; HAW; NCL; SAM |
| *Euwallacea fornicatus* (Eichhoff) | EUWAWH | 2 | Smith et al. 2019 | "Ambrosia beetles of the *Euwallacea fornicatus* (Eichhoff, 1868) species complex are emerging tree pests, responsible for significant damage to orchards and ecosystems around the world" | 24 | AFR; COM; REU; ASI; JAW; SUM; LUZ; LKA; CAM; ASI; TWN; JPN; NAM; AUS; CRL; FJI; GUM; NBR; NWG; NZL; WSM; HAW; NCL; SAM |
| *Euwallacea interjectus* (Blandford) | XYLBIN | 2 | Cognato et al 2015 | "This species has the potential to transmit tree fungal diseases given the evidence that it exacerbates fig wilt in Japanese orchards" | 13 | ASI; BOR; JAW; SUM; MEN; PHL; TWN; LKA; JPN; OKI; NAM; SAM; HAW |

**Table S-1b.** Main characteristics of the 163 Scolytinae with an invasion history. Impact and landmasses colonised

| Species | Spec code | Impact on plant health[7] | References on impact | Comments on impact | Nb of land masses | Landmasses[8] |
|---|---|---|---|---|---|---|
| *Euwallacea kuroshio* Gomez and Hulcr | EUWAKU | 2 | Smith et al. 2019 | "Ambrosia beetles of the *Euwallacea fornicatus* (Eichhoff, 1868) species complex are emerging tree pests, responsible for significant damage to orchards and ecosystems around the world" | 5 | ASI; JAW; TWN; JPN; NAM |
| *Euwallacea perbrevis (Schedl, 1951)* | EUWAPE | 2 | Smith et al. 2019 | "Ambrosia beetles of the *Euwallacea fornicatus* (Eichhoff, 1868) species complex are emerging tree pests, responsible for significant damage to orchards and ecosystems around the world" | 14 | AFR; MDG; MUS; ASI; JAW; LKA; CAM; BRB; CUB; HIS; GLP; PRI; KNA; NAM; AUS; CRL; COK; REU; WSM; HAW; SAM; SYC |
| *Euwallacea validus* (Eichhoff) | XYLBVA | 2 | Cognato et al 2015 | "*Euwallacea validus* has been implicated in the transmission of *Verticillium nonalfalfae* Inderb., a fungal disease of tree-of-heaven (*Ailanthus altissima* (Mill.) Swingle) and striped maple" | 4 | ASI; PHL; JPN; NAM |
| *Gnathotrichus materiarius* (Fitch) | GNAHMA | 0 | Inward 2019 | "*Gnathotrichus materiarius* attacks only dead and dying hosts, and damage to date has been limited to the excavation of galleries in timber and staining by the fungus" | 4 | EUR; HIS; NAM; GBR |
| *Hylastes angustatus* (Herbst) | HYASAN | 1 | Reay & Walsh 2002a | "*Hylastes angustatus* (Herbst) was reported to kill *Pinus patula* Schiede & Deppe seedlings in South Africa, but there was little quantitative assessment given (Atkinson & Govender 1997" | 3 | AFR; EUR; GBR; |
| *Hylastes ater* (Paykull) | HYASAR | 1 | Reay & Walsh 2002a | "*H. ater* attack was the dominant factor contributing to seedling mortality in the first year following planting" | 8 | AZO; ASI; JPN; AUS; EUR; GBR; NZL; SAM |
| *Hylastes linearis* Erichson | HYASLI | 0 | Mendel et al. 1997 | "Mortality due to other biotic agents, i.e., *Hylastes linearis* and *Pityophthorus pubescens* was practically nil" | 7 | AFR; CNY; MDR; EUR; GBR; COR; SAM |
| *Hylastes opacus* Erichson | HYASOP | 0 | Kumbasli et al. 2018 | "*H. opacus* breeds in the bark of stumps or at the bases of unhealthy Pinus spp." | 5 | EUR; GBR; JPN; ASI; NAM |
| *Hylastinus obscurus* (Marsham) | HYATOB | 0 | | On red clover (*Trifolium pratense*) | 7 | AFR; CNY; MDR; EUR; GBR; NAM; SAM |
| *Hylesinus toranio* (= *Bostrychus oleiperda*) (Danthione) | HYESOL | 0 | Abia et al. (2014) | On olive trees | 6 | AFR; ASI; JPN; EUR; GBR; SAM |
| *Hylurgops palliatus* (Gyllenhal) | HYLUPA | 0 | | On fallen trees | 6 | AFR; EUR; ASI; GBR; JPN; NAM |

**Table S-1b.** Main characteristics of the 163 Scolytinae with an invasion history. Impact and landmasses colonised

| Species | Spec code | Impact on plant health[7] | References on impact | Comments on impact | Nb of land masses | Landmasses[8] |
|---|---|---|---|---|---|---|
| *Hylurgus ligniperda* (Fabricius) | HYLGLI | 0 | Reay & Walsh 2002a | "there is no evidence that overwintering *H. ligniperda* larval populations are a threat to seedlings in New Zealand" | 13 | AFR; AZO; CNY; MDR; SHN; EUR; ASI; GBR; JPN; NAM; AUS; NZL; SAM |
| *Hylurgus micklitzi* Wachtl | HYLGNI | 0 | | No data on damage to plant health | 6 | EUR; SAR; SIC; ASI; AFR; MLT |
| *Hypoborus ficus* Erichson | HYPBFI | 1 | CABI 2018 | Fungi: Kolarik et al (2007). Impact: https://www.cabi.org/isc/datasheet/24078 - "*Hypoborus ficus* is one of the common pests of fig that causes drying out and death of the tree" | 7 | AFR; AZO; ASI; EUR; SAR; CNY; MDG |
| *Cryphalus scabricollis* (Eichhoff) | CRYHSC | 1 | Faccoli et al. (2016) | "Although *H. scabricollis* usually colonizes the bark of dying or stressed trees, according to what observed both in Malta and Sicily the species is able to infest also apparently healthy fig trees, killing them in a few weeks" | 8 | ASI; AND; HAI; LKA; LUZ; MLT; SIC; AFR |
| *Cryphalus dilutus* (Eichhoff) | hypdil | 0 | | No data on damage to plant health | 5 | AFR; ASI; EUR; MLT; NAM |
| *Cryphalus mangiferae* (Stebbing) | HYPCMA | 1 | Al Adawi et al. 2014 | "The mango sudden decline pathogen, *Ceratocystis manginecans*, is vectored by *Hypocryphalus mangiferae* (Coleoptera: Scolytinae) in Oman" | 22 | AFR; MDG; MUS; ASI; JAW; LKA; CAM; BRB; CUB; HIS; GLP; PRI; KNA; NAM; AUS; CRL; COK; REU; WSM; HAW; SAM; SYC |
| *Hypothenemus africanus* (Hopkins) | HYOTAF | 0 | | No data on damage to plant health | 10 | AFR; JAW; CAM; BHS; JAM; PRI; HIS; VIL; NAM; SAM |
| *Hypothenemus areccae* (Hornung) | HYOTSU | 1 | Beaver 1987 | "*H. areccae* has been recorded killing mango seedlings in Malaya" | 21 | AFR; ASI; IDN; PHL; LKA; CAM; BHS; MTQ; PRI; VIL; EUR; NAM; CRL; FJI; MRQ; MHL; NCL; AUS; NZL; HAW; SAM |
| *Hypothenemus birmanus* (Eichhoff) | HYOTBI | 1 | Beaver 1987 | "*H. birmanus* contributed to the death of mango transplants in Western Samoa" | 27 | MDG; ASI; COK; FJI; IDN; PHL; SYC; LKA; CAM; CUB; JAM; PRI; JPN; NAM; AUS; GUM; HEN; FSM; NBR; NCL; NZL; PNG; WSM; SCI; HAW; SAM; GAL |
| *Hypothenemus brunneus* Hopkins | HYOTBR | 0 | Bright & Peck 1998 | "Adults attack injured or broken branches." | 11 | CAM; BHS; CUB; GLP; JAM; PRI; VIL; NAM; SAM; TTO; GAL |
| *Hypothenemus californicus* Hopkins | HYOTCA | 0 | Bright & Peck 1998 | "This species feeds on a wide variety of host plants" | 7 | AFR; CAM; LCA; ASI; NAM; SAM; GAL |
| *Hypothenemus columbi* Hopkins | HYOTCO | 0 | Bright & Peck 1998 | "The biology of this species is similar to that of *H. eruditus*" | 9 | CAM; ATG; BHS; CUB; PRI; NAM; SAM; TTO; GAL |

**Table S-1b.** Main characteristics of the 163 Scolytinae with an invasion history. Impact and landmasses colonised

| Species | Spec code | Impact on plant health[7] | References on impact | Comments on impact | Nb of land masses | Landmasses[8] |
|---|---|---|---|---|---|---|
| *Hypothenemus crudiae* (Panzer) | HYOTHI | 2 | Wood 1977; Bright & Peck 1998 | Wood 1977: "its greatest populations occur in seeds, pods, or other fruiting bodies, where it has caused much economic damage in mature seeds both in the field and in storage. Its importance in agriculture or forestry is limited to its effect on seed production." | 21 | AFR; CPV; MDG; ASI; JAW; LKA; CAM; CUB; GRD; MSR; PRI; HIS; NAM; AUS; COK; MRQ; FSM; HAW; SAM; GAL; TTO; |
| *Hypothenemus elephas* (Eichhoff) | hypele | 0 | | No data on damage to plant health | 4 | AFR; MDG; MUS; SAM |
| *Hypothenemus erectus* LeConte | HYOTER | 0 | | No data on damage to plant health | 7 | CAM; BHS; CUB; JAM; VIL; NAM; SAM |
| *Hypothenemus eruditus* Westwood | HYOTEU | 1 | Kambestad et al 2017 | "*H. eruditus* usually breeds in dead, sometimes quite dry, host tissues, and is generally not considered to be a pest. However, attacks on commercially important plants such as coffee, cocoa, macadamia nuts, camphor trees, plane trees, and timber of rubberwood have been reported" | 37 | AFR; AZO; CNY; MDG; SYC; ASI; JAW; SUM; MRQ; PHL; LKA; CAM; BHS; CUB; DMA; GLP; JAM; NEV; PRI; HIS; VCT; VIL; GBR; EUR; TWN; JPN; NAM; AUS; COK; FJI; FSM; NCL; HAW; SAM; GAL; TTO; MLT |
| *Hypothenemus hampei* (Ferrari) | STEHHA | 2 | Vega et al. 2015 | on coffee berries | 18 | AFR; CNY; FER; STP; ASI; JAW; SUM; PHL; LKA; CAM; JAM; PRI; NAM; FSM; NCL; WSM; TAH; SAM |
| *Hypothenemus javanus* (Eggers) | HYOTJA | 0 | Vega et al. 2015 | | 17 | AFR; STP; ASI; JAW; PHL; LKA; CAM; BHS; CUB; P; GLHIS; MTQ; TWN; NAM; HAW; SAM; SYC |
| *Hypothenemus leprieuri* ( Perris) | HYOTLE | 0 | | No data on damage to plant health | 5 | AFR; ASI; SAR; CYP; MLT |
| *Hypothenemus obscurus* (Fabricius) | HYOTOB | 1 | Vega et al. 2015 | "...the second most economically damaging species in the genus (after the coffee berry borer)" | 12 | JAW; CAM; CUB; GLP; JAM; PRI; HIS; NAM; HAW; SAM; GAL; TTO |
| *Hypothenemus plumeriae* (Nordlinger) | STEHPL | 0 | | No data on damage to plant health | 9 | AFR; CAM; GLP; HIS; NAM; HAW; SAM; GAL; TTO |
| *Hypothenemus pubescens* Hopkins | HYOTPB | 0 | | No data on damage to plant health | 4 | PRI; NAM; HAW; SAM |
| *Hypothenemus seriatus* (Eichhoff) | STEHSE | 1 | Vega et al. 2015 | "This species is also known to damage cocoa seedlings" | 22 | AFR; MDG; SYC; JAW; PHL; LKA; CAM; BHS; BRB; CUB; HIS; PRI; DMA; VIL; ASI; NAM; AUS; FJI; FSM; HAW; SAM; GAL |
| *Hypothenemus setosus* (Eichhoff) | HYOTSE | 0 | | No data on damage to plant health | 11 | AFR; MDG; CAM; CUB; GLP; HIS; JAM; PRI; TWN; NAM; SAM |
| *Ips calligraphus* (Germar) | IPSXCA | 0 | | No data on damage to plant health | 5 | LUZ; BHS; JAM; HIS; NAM; |

**Table S-1b.** Main characteristics of the 163 Scolytinae with an invasion history. Impact and landmasses colonised

| Species | Spec code | Impact on plant health[7] | References on impact | Comments on impact | Nb of land masses | Landmasses[8] |
|---|---|---|---|---|---|---|
| *Ips cembrae* (Herr) | IPSXCE | 1 | | Fungi: Redfern, D. B., Stoakley, J. T., Steele, H., & Minter, D. W. (1987). Dieback and death of larch caused by *Ceratocystis laricicola* sp. nov. following attack by *Ips cembrae*. Plant Pathology, 36(4), 467-480. | 2 | EUR; GBR; |
| *Ips grandicollis* (Eichhoff) | IPSXGR | 1 | CABI 2018 | Fungi: Zhou et al. 2007; Stone & Simpson, 1989. Impact: https://www.cabi.org/isc/datasheet/28825 | 5 | BHS; HIS; JAM; AUS; NAM |
| *Kissophagus hederae* (Schmidt) | KISSHE | 0 | | No data on damage to plant health | 7 | AFR; EUR; SAR; SIC; GBR; MLT; ASI |
| *Liparthrum artemisiae* Wollaston | lipart | 0 | | No data on damage to plant health | 2 | MDR; CNY |
| *Liparthrum bituberculatum* Wollaston | lipbit | 0 | | No data on damage to plant health | 2 | MDR; CNY |
| *Liparthrum curtum* Wollaston | lipcur | 0 | | No data on damage to plant health | 3 | AZO; MDR; CNY |
| *Liparthrum inarmatum* Wollaston | lipina | 0 | | No data on damage to plant health | 3 | MDR; CNY; AFR |
| *Liparthrum mandibulare* Wollaston | lipman | 0 | | No data on damage to plant health | 2 | MDR; CNY |
| *Liparthrum mori* (Aube) | LPRTMO | 0 | | No data on damage to plant health | 4 | AFR; EUR; SAR; MLT |
| *Microborus boops* Blandford | micboop | 0 | Wood 1977 | Branches | 6 | AFR; MDG; CAM; JAM; SAM; TTO |
| *Microperus* (*Coptodryas*) *eucalypticus* (Schedl) | MIPREU | 0 | Beaver & Liu 2016 | "(…) with the exception of *M. eucalypticus*, (…) all have been recorded as pest species" | 3 | AUS; NZL; NCL |
| *Microperus quercicola* (Eggers) | micquer | 0 | | No data on damage to plant health | 3 | ASI; JPN; TWN |
| *Monarthrum mali* (Fitch) | MNTHMA | 0 | Kirkendall et al. 2008 | "The eastern USA species *M. mali* and *M. fasciatum* attack dying, injured, or recently cut trees" | 8 | EUR; BHS; CUB; DMA; HIS; GLP; PRI; NAM |
| *Orthotomicus angulatus* (Eichhoff) | ortang | 0 | | No data on damage to plant health | 3 | ASI; JPN; TWN |
| *Orthotomicus caelatus* (Eichhoff) | ORTCCA | 0 | | No data on damage to plant health | 3 | AFR; BHS; NAM |
| *Orthotomicus erosus* (Wollaston) | IPSXER | 1 | CABI 2018; Mendel 1987 | https://www.cabi.org/isc/datasheet/37954; Mendel 1987: "Severe outbreaks occur after excessive thinning followed by a winter with low rainfall, in plots with poor phytosanitation, where there is a lack of ecological adaption of pine species, or after fire in adjoining stands. P. *calcaratus* also attacks young seedlings affected by the Israeli pine bast scale." | 13 | EUR; AZO; COR; GBR; SAR; SIC; MLT; AFR; MDR; FJI; ASI; NAM; SAM |
| *Orthotomicus laricis* (Fabricius) | IPSXLC | 0 | | Expert knowledge | 9 | AFR; ASI; JPN; SAK; EUR; COR; GBR; SAR; SAM |
| *Orthotomicus proximus* (Eichhoff) | IPSXPR | 0 | | No data on damage to plant health | 4 | ASI; JPN; EUR; COR; MDG |

**Table S-1b.** Main characteristics of the 163 Scolytinae with an invasion history. Impact and landmasses colonised

| Species | Spec code | Impact on plant health[7] | References on impact | Comments on impact | Nb of land masses | Landmasses[8] |
|---|---|---|---|---|---|---|
| *Pagiocerus frontalis* (Fabricius) | PAGIFR | 0 | Kirkendall et al. 2015 | "often collected from seeds of Lauraceae, including commercial avocado (*Persea americana* Mill.). In Mexico, it bores into partially or completely exposed seeds lying on the ground and does not attack fruits on the tree" | 8 | GAL; NAM; CUB; GLP; HIS; JAM; CAM; SAM |
| *Phloeosinus armatus* Reitter | PHLSAR | 1 | Mendel 1984 | "They breed on stems and branches of well-developed trees which are suffering from sudden stress due to drought, fire, root damage, or heavy infestation by the fungus *Seiridium cardinale* (Wag.)" | 4 | CYP; EUR; ASI; NAM |
| *Phloeosinus cupressi* Hopkins | PHLSCU | 0 | | https://www.nzffa.org.nz/farm-forestry-model/the-essentials/forest-health-pests-and-diseases/Pests/Phloeosinus-cupressi/the-cypress-bark-beetle/: "The cypress barkbeetle breeds under the bark of dead and dying trees but is more often found under the bark of felled trees or in dead branches." | 4 | CAM; NAM; AUS; NZL |
| *Phloeosinus rudis* Blandford | PHLSRD | 1 | Moraal 2010; Moucheron et al. 2019 | On weakened trees. "In the summer of 2004, hundreds of shrubs and trees of Cupressaceae in The Netherlands were killed by the Japanese cypress bark beetle, *Phloeosinus rudis*." | 3 | ASI; JPN; EUR |
| *Phloeosinus thujae* (Perris) | PHLSTH | 1 | Moraal 2010 | "In addition [to P. rudis], the Mediterranean cypress bark beetles *Phloeosinus bicolor* and *Phloeosinus thujae* were identified as well as the cause of death of many Cupressaceae on several locations in 2004." | 9 | AFR; ASI; MLT; EUR; GBR; COR; SAR; SIC; CNY |
| *Phloeotribus liminaris* (Harris) | PHLBLI | 1 | Pennacchio et al 2004 | "The beetle's wintering activity is particularly harmful, as it digs refuges in the vital internal bark of trees in good vegetative condition" | 2 | NAM; EUR |
| *Phloeotribus scarabaeoides* (Bernard) | PHLBOL | 1 | González & Campos (1994) | No data on damage to plant health | 6 | AFR; ASI; EUR; CYP; COR; SAR |
| *Pityogenes bidentatus* (Herbst) | PITYBD | 0 | | No data on damage to plant health | 5 | MDG; EUR; GBR; ASI; NAM |

**Table S-1b.** Main characteristics of the 163 Scolytinae with an invasion history. Impact and landmasses colonised

| Species | Spec code | Impact on plant health[7] | References on impact | Comments on impact | Nb of land masses | Landmasses[8] |
|---|---|---|---|---|---|---|
| *Pityogenes calcaratus* (Eichhoff) | PITYCC | 1 | Mendel 1987 | "Severe outbreaks occur after excessive thinning followed by a winter with low rainfall, in plots with poor phytosanitation, where there is a lack of ecological adaption of pine species, or after fire in adjoining stands. P. calcaratus also attacks young seedlings affected by the Israeli pine bast scale." | 6 | MLT; EUR; COR; SAR; AFR; ASI |
| *Pityogenes chalcographus* (Linnaeus) | PITYCH | 1 | Hedgren 2004 | "this study confirms *P. chalcographus* as a species restricted to weakened, or recently killed trees" | 7 | JAM; ASI; KUR; SAK; JPN; EUR; GBR |
| *Pityokteines curvidens* (Germar) | PITKCU | 1 | CABI 2018 | https://www.cabi.org/isc/datasheet/45720 | 4 | AFR; ASI; JPN; EUR |
| *Pityophthorus juglandis* Blackman | PITOJU | 2 | EPPO 2015 | Fungi: Kolarik et al 2011; pheromone: not in Pherobase but described in a patent by Seybold & USDA | 2 | EUR; NAM |
| *Pityophthorus solus* (Blackman) | PITOSO | 0 | | No data on damage to plant health | 2 | NAM; EUR |
| *Planiculus (Euwallacea) bicolor* (Blandford) | euwabic | 0 | Beaver 1988 | No data on damage to plant health | 11 | ASI; SYC; AND; JPN; LKA; FJI; BOR; JAW; LUZ; WSM; SLB |
| *Polygraphus poligraphus* (Linnaeus) | POLGPO | 0 | | No data on damage to plant health | 6 | AFR; JPN; SAK; ASI; EUR; GBR; |
| *Polygraphus proximus* Blandford | POLGPR | 1 | Krivets et al. 2015 | "this species is currently one of the main factors of degradation of the Siberian fir forests " | 4 | ASI; JPN; SAK; EUR |
| *Polygraphus rufipennis* (Kirby) | POLGRU | 0 | Bowers et al. 1996 | No data on damage to plant health | 2 | NAM; AFR |
| *Premnobius ambitiosus* (Schaufuss) | preamb | 0 | | No data on damage to plant health | 4 | AFR; CNY; CAM; SAM |
| *Premnobius cavipennis* Eichhoff | PREBCA | 0 | Rabaglia et al. 2006 | No data on damage to plant health | 11 | AFR; MDG; CAM; BHS; CUB; DMA; JAM; PRI; NAM; SAM; TTO |
| *Pseudohylesinus sericeus* (Mannerheim) | PSDHSE | 0 | | No data on damage to plant health | 2 | JPN; NAM |
| *Scolytogenes jalapae (Letzner)* | scojal | 0 | | No data on damage to plant health | 10 | CAM; CUB; GLP; JAM; HIS; PRI; VIL; JPN; NAM; SAM |
| *Scolytoplatypus tycon* Blandford | scotyc | 0 | | No data on damage to plant health | 4 | ASI; JPN; SAK; EUR |
| *Scolytus amygdali* Guerin-Meneville | SCOLAM | 0 | Zeiri et al. 2018 | These beetles target weak, old trees, where they make galleries and holes in the bark. | 9 | EUR; COR; SAR; SIC; MLT; CNY; AFR; CYP; ASI |
| *Scolytus dimidiatus* Chapuis | scodim | 0 | | No data on damage to plant health | 5 | CAM; CUB; JAM; NAM; SAM |

**Table S-1b.** Main characteristics of the 163 Scolytinae with an invasion history. Impact and landmasses colonised

| Species | Spec code | Impact on plant health[7] | References on impact | Comments on impact | Nb of land masses | Landmasses[8] |
|---|---|---|---|---|---|---|
| *Scolytus kirschi* Skahtzky | SCOLKI | 2 | Six et al. 2005 | Fungi: Six et al. 2005. Impact: Six et al. 2005: "*Scolytus kirschii* is a serious pest of elms, capable of killing healthy trees, resulting in considerable economic impact. Furthermore, the beetle is capable of being the vector of the pathogens responsible for Dutch elm disease (DED), *Ophiostoma ulmi* and *Ophiostoma novo-ulmi*. | 4 | ASI; AFR; EUR; SIC |
| *Scolytus mali* (Bechstein) | SCOLMA | 0 | | No data on damage to plant health | 4 | AFR; EUR; GBR; NAM; |
| *Scolytus multistriatus* (Marsham) | SCOLMU | 2 | CABI 2018 | https://www.cabi.org/isc/datasheet/49212 | 13 | AFR; ASI; IDN; EUR; GBR; IRL; SIC; SAR; NAM; AUS; NWG; NZL; SAM |
| *Scolytus rugulosus* (Muller) | SCOLRU | 1 | | Blackman 1922 | 11 | EUR; COR; GBR; SAR; SIC; MLT; AFR; ASI; CYP; NAM; SAM |
| *Scolytus schevyrewi* Semenov | SCOLSH | 2 | CABI 2018; Jacobi et al 2013 | Fungi: Jacobi et al 2013. Impact: https://www.cabi.org/isc/datasheet/49200; "The inoculation of trees via feeding wounds was successful 30% of the time for in-vivo trials and 33% for in-vitro trials" | 2 | ASI; NAM |
| *Scolytus sulcifrons* Rey | SCOLSU | 0 | | No data on damage to plant health | 3 | EUR; ASI; MLT |
| *Thamnurgus characiae* Rosenhauer | THMNCH | 0 | | No data on damage to plant health | 5 | EUR; SAR; SIC; MLT; AFR |
| *Theoborus ricini* (Eggers) | theori | 0 | | No data on damage to plant health | 7 | AFR; STP; CAM; PRI; HIS; NAM; SAM |
| *Tomicus piniperda* (Linnaeus) | BLASPI | 1 | Langstrom and Hellqvist 1993 | Shoot cutting, sometimes massive | 9 | AFR; MDR; ASI; PHL; JPN; TWN; EUR; GBR; NAM |
| *Truncaudum* (*Cyclorhipidion*) *agnatum* (Eggers) | TRUCAG | 0 | Sanguansub et al 2012 | Truncaudum agnatum collected from cut logs | 11 | ASI; NCL; AUS; BIS; BOR; JAW; CRL; PLW; NWG; LUZ; SLB |
| *Trypodendron domesticum* (Linnaeus) | TRYDDO | 1 | La Spina et al. 2013 | | 6 | EUR; GBR; IRL; SIC; SAR; NAM |
| *Xyleborinus andrewesi* (Blandford) | XYBIAN | 0 | Okins & Thomas 2009 | "Unless they occur in very large numbers, damage is minimal (...) *Xyleborinus andrewesi* is unlikely to become even a minor pest" | 14 | AFR; SYC; ASI; IDN; JAW; PHL; LKA; CUB; JAM; JPN; NAM; MAR; NWG; HAW |
| *Xyleborinus artestriatus* (Eichhoff) | XYBIAR | 0 | Gomez et al. 2018 | | 7 | ASI; IDN; LKA; NAM; AUS; FJI; NWG |

**Table S-1b.** Main characteristics of the 163 Scolytinae with an invasion history. Impact and landmasses colonised

| Species | Spec code | Impact on plant health[7] | References on impact | Comments on impact | Nb of land masses | Landmasses[8] |
|---|---|---|---|---|---|---|
| *Xyleborinus attenuatus* (Blandford) | XYBIAL | 0 | SLU 2017 | "There is however little support in the easily accessible literature that the pest cause significant damage. This is in agreement with a previous literature search that showed that there had not been a single report of damage by *X. attenuatus* on living trees in USA" | 4 | ASI; JPN; EUR; NAM |
| *Xyleborinus exiguus* (Walker) | XYBIEX | 0 | | No data on damage to plant health | 20 | AFR; ASI; IDN; JAW; SUM; PHL; LKA; CAM; AUS; COK; FJI; GUM; MAR; NCL; NUE; PLW; PNG; WSM; SCI; SLB |
| *Xyleborinus gracilis* Eichhoff | XYBIGR | 0 | Rabaglia et al. 2006 | | 6 | GAL, NAM; AZO; CAM; GLP; SAM |
| *Xyleborinus octiesdentatus* (Murayama) | XYBIOC | 0 | | No data on damage to plant health | 3 | ASI; JPN; NAM |
| *Xyleborinus saxeseni* (Ratzeburg) | XYLBSA | 2 | Rabaglia et al. 2006 | "It is one of the most damaging and occasionally aggressive species in the tribe in North America." | 24 | EUR; AZO; COR; GBR; SAR; SIC; MLT; AFR; CNY; MDR; ASI; JPN; PHL; TWN; NAM; GUM; NWG; AUS; NZL; WSM; HAW; SAM; GAL; NCL |
| *Xyleborus affinis* Eichhoff | XYLBAF | 1 | Rabaglia et al. 2006 | "As with *X. ferrugineus*, this species can cause economic damage in moist, lowland areas of the Neotropics." | 33 | AFR; AZO; FER; MDG; MUS; REU; SYC; IDN; LKA; CAM; BRB; CUB; DMA; GLP; JAM; PRI; HIS; VCT; ASI; NAM; AUS; COK; FJI; MAR; NCL; PLW; WSM; TAH; HAW; SAM; GAL; TTO; EUR |
| *Xyleborus africanus* Eggers | xylafr | 0 | | No data | 2 | AFR; SYC |
| *Xyleborus atratus* Eichhoff | xylatr | 0 | | No data on damage to plant health | 7 | ASI; JPN; TWN; PHL; IDN; NWG; NAM; EUR |
| *Xyleborus bispinatus* Eichhoff | XYLBBI | 0 | Faccoli et al. (2016) | "*X. bispinatus* (…) was found in Sicily only in dying or recently killed fig trees both in 2014 and 2015 " | 7 | CAM; HIS; NAM; PNG; SAM; TTO; EUR |
| *Xyleborus ferrugineus* (Fabricius) | XYLBFE | 2 | Rabaglia et al. 2006 | "As with *X. ferrugineus*, this species can cause economic damage in moist, lowland areas of the Neotropics." | 31 | AFR; AZO; CPV; FER; MDG; REU; SYC; CAM; BHS; CUB; DMA; GLP; JAM; PRI; HIS; VIL; NAM; WSM; AUS; COK; FJI; GUM; MRQ; NCL; NWG; TAH; TKL; HAW; SAM; GAL; TTO |

**Table S-1b.** Main characteristics of the 163 Scolytinae with an invasion history. Impact and landmasses colonised

| Species | Spec code | Impact on plant health[7] | References on impact | Comments on impact | Nb of land masses | Landmasses[8] |
|---|---|---|---|---|---|---|
| *Xyleborus glabratus* Eichhoff | XYLBGR | 2 | Hughes et al. 2017 | USDA Forest Service Forest Inventory and Analysis data were used to estimate that over 300 million trees of redbay (*Persea borbonia sensu lato*) have succumbed to the disease since the early 2000s (ca 1/3 of the pre-invasion population) | 4 | ASI; JPN; TWN; NAM |
| *Xyleborus monographus* (Fabricius) | XYLBMO | 0 | | No data on damage to plant health | 4 | AFR; EUR; GBR; NAM |
| *Xyleborus perforans* (Wollaston) | XYLBPE | 1 | CABI 2018 | https://www.cabi.org/isc/datasheet/57169 - X. *perforans* has been recorded as a minor pest of sugarcane and coconut trees in Indonesia (Kalshoven, 1964; Browne, 1968), and in the days of wooden beer, wine and rum barrels, was known to bore into casks and cause leakage (Blandford, 1893; Schedl, 1963). It has been known to cause minor damage by its attacks on the tapped panels of rubber trees in Guyana and Sri Lanka (Browne, 1968), and on coffee and coffee shade trees in Suriname (LePelley, 1968). However, it is more important in many areas because of its heavy attacks on newly felled trees and recently sawn, unseasoned timber. The attacks result in numerous pinholes in the wood and fungal staining around them (Browne, 1961), and can render the timber unusable for furniture or veneer. " | 36 | AFR; ASI; NAM; CPV; EUR; FJI; IDN; JPN; MDG; PHL; SLB; LKA; TWN; TON; BIS; AUS; AZO; CNY; COK; AZO; GUM; KIR; MDR; MAR; MUS; FSM; NCL; NWG; VAN; NUE; OKI; PLW; SYC; SCI; TAH; TUA |
| *Xyleborus pfeilii* (Ratzeburg) | XYLBPF | 0 | CDFA 2018 | https://blogs.cdfa.ca.gov/Section3162/?p=5255 | 7 | AFR; EUR; ASI; JPN; NAM; NZL; SAM |
| *Xyleborus seriatus* Blandford | XYLBSE | 0 | EPPO Reporting Service 2006-10 | Recent finding and so far, no reports of hosts, damage or expansion in USA. In Asia, many conifer and hardwood species are reported as hosts (*Acer, Aesculus, Betula Chamaecyparis, Cryptomeria, Fagus, Larix, Pinus, Prunus, Quercus, Thuja, Tilia* and *Tsuga*). | 3 | ASI; JPN; NAM |
| *Xyleborus spinulosus* Blandford | XYLBSN | 0 | Gomez et al. 2018 | | 6 | CAM; GAL; HIS; VIL; NAM; HAW; |

**Table S-1b.** Main characteristics of the 163 Scolytinae with an invasion history. Impact and landmasses colonised

| Species | Spec code | Impact on plant health[7] | References on impact | Comments on impact | Nb of land masses | Landmasses[8] |
|---|---|---|---|---|---|---|
| *Xyleborus volvulus* (Fabricius) | xylvol | 0 | Menocal et al. 2017 | "Although *X. volvulus* has not been associated with economic damage to trees..." | 25 | AFR; MUS; MDG; SYC; ASI; TWN; PHL; CAM; SAM; GAL; ATG; BHS; CUB; GRD; JAM; HIS; VIL; JPN; NAM; WSM; AUS; NCL; HAW; TTO; SLB |
| *Xylosandrus* (*Apoxyleborus*) *mancus* (Blandford) | xylman | 0 | | No data on damage to plant health | 10 | AFR; SYC; ASI; JPN; TWN; JAW; SUM; LKA; MDG; PHL |
| *Xylosandrus amputatus* (Blandford) | XYLSAM | 0 | Bateman et al. 2015 | "Since it was first detected in Florida, this species has spread north to Georgia, but has not yet been found attacking healthy trees nor causing economic damages in the United States." | 4 | ASI; JPN; TWN; NAM |
| *Xylosandrus compactus* (Eichhoff) | XYLSCO | 2 | CABI 2019 | https://www.cabi.org/isc/datasheet/57234 | 27 | AFR; COM; FER; MDG; REU; SYC; ASI; IDN; PHL; LKA; VIL; CUB; DMA; PRI; TWN; JPN; NAM; FJI; GUM; NZL; WSM; HAW; NCL; SAM; TTO; EUR; SIC |
| *Xylosandrus crassiusculus* (Motschulsky) | XYLSCR | 2 | CABI 2019 | https://www.cabi.org/isc/datasheet/57235 | 25 | AFR; FER; MDG; MUS; SYC; ASI; IDN; SUM; PHL; LKA; CAM; PRI; TWN; JPN; EUR; NAM; GUM; NCL; PLW; PNG; WSM; HAW; AUS; SAM; NZL |
| *Xylosandrus germanus* (Blandford) | XYLSGE | 1 | Galko et al. 2019 | While living but weakened trees in Europe and North America are attacked by *X. germanus*, the greatest negative impact within Slovakia is attacks on recently felled logs of oak, beech and spruce trees, which provide high quality timber/lumber. | 7 | ASI; EUR; GBR; JPN; TWN; NAM; HAW |
| *Xylosandrus morigerus* (Blandford) | XYLSMO | 2 | Beaver 1987 | "*X. morigerus* attacked tree seedlings, including tea and mahogany in Java (Kalshoven 1961). It is also a potential pest of established living trees as a twig borer. It is an important pest of coffee in several countries and often attacks shade and ornamental trees" | 25 | MDG; ASI; IDN; MUS; PHL; LKA; CAM; PRI; GBR; EUR; TWN; NAM; AUS; CRL; FJI; GUM; MAR; NBR; NCL; PNG; WSM; HAW; SAM; GAL; SYC |
| *Xylosandrus pseudosolidus* (Schedl) | XYLSPS | 0 | Brockerhoff et al. (2003) | | 3 | AUS; TAS; NZL |
| *Xyloterinus politus* (Say) | xypol | 0 | | No data on damage to plant health | 2 | NAM; EUR |

**Table S-1b.** Main characteristics of the 163 Scolytinae with an invasion history. Impact and landmasses colonised

### Table S-1c. References


Abia, M. M. C., Alarcón, J. C., & Belenguer, M. S. N. (2014). El barrenillo del olivo *Hylesinus toranio* (Coleoptera, Curculionidae, Scolytinae) Una plaga importante del campo aragonés. *Phytoma España: La revista profesional de sanidad vegetal*, (260), 48-52.

Al Adawi, A. O., Al Jabri, R. M., Deadman, M. L., Barnes, I., Wingfield, B., & Wingfield, M. J. (2013). The mango sudden decline pathogen, *Ceratocystis manginecans*, is vectored by *Hypocryphalus mangiferae* (Coleoptera: Scolytinae) in Oman. *European journal of plant pathology*, *135*(2), 243-251.

Atkinson TH, Rabaglia RJ, Bright DE (1990) Newly detected exotic species of *Xyleborus* (Coleoptera: Scolytidae) with a revised key to species in eastern North America. *The Canadian Entomologist* 122: 92 – 104. https://doi.org/10.4039/Ent12293-1

Atkinson, TH. 2021. Bark and Ambrosia Beetles. http://www.barkbeetles.info/index.php. Last accessed on 10 February 2021.

Bateman, C., Kendra, P. E., Rabaglia, R., & Hulcr, J. (2015). Fungal symbionts in three exotic ambrosia beetles, *Xylosandrus amputatus*, *Xyleborinus andrewesi*, and *Dryoxylon onoharaense* (Coleoptera: Curculionidae: Scolytinae: Xyleborini) in Florida. *Symbiosis*, *66*(3), 141-148.

Beaver, R. A. (1976). The biology of Samoan bark and ambrosia beetles (Coleoptera, Scolytidae and Platypodidae). *Bulletin of Entomological Research*, *65*(4), 531-548.

Beaver, R. A. (1987). Biological studies on bark beetles of the Seychelles (Col., Scolytidae and Platypodidae). *Journal of Applied Entomology*, 104 (1)5), 11-23

Beaver, R. A. (1988). Biological studies on ambrosia beetles of the Seychelles (Col., Scolytidae and Platypodidae). *Journal of Applied Entomology*, *105* (1-5), 62–73.

Beaver, R. A., & Gebhardt, H. (2006). A review of the oriental species of Scolytoplatypus Schaufuss (Coleoptera, Curculionidae, Scolytinae). *Deutsche Entomologische Zeitschrift*, *53*(2), 155-178.

Beaver, R. A., & Liu, L. Y. (2016). Ambrosia beetles (Coleoptera: Curculionidae, Scolytinae, Xyleborini) of New Caledonia, with descriptions of four new species. *Entomologist's Monthly Magazine*, *152*, 21–32.

Bevan, D., & King, C. J. (1983). *Dendroctonus micans* Kug, - a new pest of spruce in UK. *The Commonwealth Forestry Review*, 41-51.

Blackman, M.W., 1922. *Mississippi bark beetles*. Miss., Agric. Exp. Stn.,Tech. Bull, 11,130 pp., 18 pls.

Bowers, W. W., Borden, J. H., & Raske, A. (1996). Incidence and impact of *Polygraphus rufipennis* (Coleoptera: Scolytidae) in Newfoundland. *Forest ecology and management*, *89*(1-3), 173-187.

Bright, D. E. 2021. Catalog of Scolytidae (Coleoptera), supplement 4 (2011-2019) with an annotated checklist of the world fauna (Coleoptera: Curculionoidea: Scolytidae), A (Doctoral dissertation, Colorado State University. Libraries). https://mountainscholar.org/handle/10217/229307

Bright, D. E., & Peck, S. B. (1998). Scolytidae from the Galápagos Islands, Ecuador, with descriptions of four new species, new distribution records, and a key to species (Coleoptera: Scolytidae). *Koleopterologische Rundschau*, *68*, 233-252.

Brockerhoff, E. G., & Bain, J. (2000). Biosecurity implications of exotic beetles attacking trees and shrubs in New Zealand. *New Zealand Plant Protection*, *53*, 321-327.

Brockerhoff, E. G., Knížek, M., & Bain, J. (2003). Checklist of indigenous and adventive bark and ambrosia beetles (Curculionidae: Scolytinae and Platypodinae) of New Zealand and interceptions of exotic species (1952-2000). *New Zealand Entomologist*, *26*(1), 29-44.

Brockerhoff E.G., Bain J., Kimberley M., Knížek M. 2006. Interception frequency of exotic bark and ambrosia beetles (Coleoptera: Scolytinae) and relationship with establishment in New Zealand and worldwide. *Canadian Journal of Forest Research*, 36 (2): 289–298.

CABI 2021a. CABI Invasive species compendium. Datasheet on *Cryphalus pallidus*. https://www.cabi.org/isc/datasheet/113633. Consulted on 15 February 2021.

CABI 2021b. CABI Invasive species compendium. Datasheet on *Anisandrus dispar*. https://www.cabi.org/isc/datasheet/57157. Consulted on 15 February 2021.

CABI 2021c. CABI Invasive species compendium. Datasheet on *Xylosandrus mutilatus*. https://www.cabi.org/isc/datasheet/57239. Consulted on 15 February 2021.

CABI 2021d. CABI Invasive species compendium. Datasheet on *Euwallacea piceus*. https://www.cabi.org/isc/datasheet/57171. Consulted on 15 February 2021.

CABI 2021e. CABI Invasive species compendium. Datasheet on *Hypoborus ficus*. https://www.cabi.org/isc/datasheet/24078. Consulted on 15 February 2021.

CABI 2021f. CABI Invasive species compendium. Datasheet on *Ips grandicollis*. https://www.cabi.org/isc/datasheet/28825. Consulted on 15 February 2021.

CABI 2021g. CABI Invasive species compendium. Datasheet on *Pityokteines curvidens*. https://www.cabi.org/isc/datasheet/45720. Consulted on 15 February 2021.

CABI 2021h. CABI Invasive species compendium. Datasheet on *Scolytus multistriatus*. https://www.cabi.org/isc/datasheet/49212 Consulted on 15 February 2021.

CABI 2021i. CABI Invasive species compendium. Datasheet on *Scolytus schevyrewi*. https://www.cabi.org/isc/datasheet/49200. Consulted on 15 February 2021.

CABI 2021j. CABI Invasive species compendium. Datasheet on *Xyleborus perforans*. https://www.cabi.org/isc/datasheet/57169. Consulted on 15 February 2021.

CABI 2021k. CABI Invasive species compendium. Datasheet on *Xyleborus similis*. https://www.cabi.org/isc/datasheet/57175. Consulted on 15 February 2021.

CABI 2021l. CABI Invasive species compendium. Datasheet on *Xylosandrus compactus*. https://www.cabi.org/isc/datasheet/57234. Consulted on 15 February 2021.

### Table S-1c. References


CABI 2021m. CABI Invasive species compendium. Datasheet on *Xylosandrus crassiusculus.* https://www.cabi.org/isc/datasheet/57235. Consulted on 15 February 2021.

Cognato AI, Hoebeke ER, Kajimura H, Smith SM (2015) History of the Exotic Ambrosia Beetles *Euwallacea interjectus* and *Euwallacea validus* (Coleoptera : Curculionidae: Xyleborini) in the United States. *Journal of Economic Entomology* 108 (3): 1129–1135. https://doi.org/10.1093/jee/tov073

Cognato, A. I., R. O'Donnell-Olson & R. J. Rabaglia. 2011. A native Asian ambrosia beetle *Xylosandrus amputatus* (Blandford) (Curculionidae: Scolytinae: Xyleborini) discovered in Florida, U.S.A. *Coleopterists Bulletin* 65:43 – 45.

Cognato, A. I., Rabaglia, R. J., & Vandenberg, N. J. (2013). Another Asian ambrosia beetle, *Xyleborinus artestriatus* (Eichhoff 1878) (Coleoptera: Curculionidae: Scolytinae: Xyleborini), newly detected in North America. *The Pan-Pacific Entomologist*, 89(1), 27–31.

Coleman, T.W., Poloni, A.L., Chen, Y. *et al.* Hardwood injury and mortality associated with two shot hole borers, *Euwallacea* spp., in the invaded region of southern California, USA, and the native region of Southeast Asia. *Annals of Forest Science* **76,** 61 (2019). https://doi-org.ezproxy.ulb.ac.be/10.1007/s13595-019-0847-6

Cooperband, M. F., Cossé, A. A., Jones, T. H., Carrillo, D., Cleary, K., Canlas, I., & Stouthamer, R. (2017). Pheromones of three ambrosia beetles in the *Euwallacea fornicatus* species complex: ratios and preferences. *PeerJ*, *5*, e3957.

Crooke, M. & Bevan, D. (1957). Note on the first British occurrence of *Ips cembrae* Heer (Col. Scolytidae). *Forestry*, *30*(1), 21-28.

Dodelin, B., Saurat, R. (2017) Espèce invasive nouveau pour la faune de France Scolyte. *Xyloterinus politus* a traversé l'Atlantique. https://entomodata.wordpress.com/2017/07/08/xyloterinus-politus-a-traverse-latlantique/. Accessed on 4 December 2020.

EPPO GD. EPPO Global Database, https://gd.eppo.int/. Last consulted on 13 February 2021.

EPPO. 2013. First report of *Anisandrus maiche* in Ukraine. *EPPO Reporting Service* no. 02–2013 Num. article: 2013/030

EPPO. 2019. First report of *Xylosandrus crassiusculus* in New Zealand. *EPPO Reporting Service* no. 04–2019. Num. article: 2019/074.

EPPO 2020a. EPPO Datasheet: *Pityophthorus juglandis*. https://gd.eppo.int/taxon/PITOJU/datasheet. Last consulted on 15 February 2021.

EPPO 2006. Exotic bark- and wood-boring Coleoptera in the USA. EPPO Reporting Service no. 10 – 2006. Num. article: 2006/208. https://gd.eppo.int/reporting/article-1278. Last consulted on 21 February 2021.

Faccoli, M. (2008). First record of *Xyleborus atratus* Eichhoff from Europe, with an illustrated key to the European Xyleborini (Coleoptera: Curculionidae: Scolytinae). *Zootaxa*, 1772(1), 55-62

Faccoli, M., Campo, G., Perrotta, G., & Rassati, D. 2016. Two newly introduced tropical bark and ambrosia beetles (Coleoptera: Curculionidae, Scolytinae) damaging figs (*Ficus carica*) in southern Italy. *Zootaxa*, 4138 (1), 189–194.

Faccoli, M., Frigimelica, G., Mori, N., Petrucco Toffolo, E., Vettorazzo, M., & Simonato, M. (2009). First record of *Ambrosiodmus* (Hopkins, 1915)(Coleoptera: Curculionidae, Scolytinae) in Europe. *Zootaxa*, *2303*(1), 57-60

Faccoli, M., Simonato, M., & Petrucco Toffolo, E. (2012). First record of *Cyrtogenius* Strohmeyer in Europe, with a key to the European genera of the tribe Dryocoetini (Coleoptera: Curculionidae, Scolytinae). *Zootaxa*, *3423* (1), 27–35.

Farm Forestry New Zealand. 2021. Pests and diseases of forestry in New Zealand. https://www.nzffa.org.nz/farm-forestry-model/the-essentials/forest-health-pests-and-diseases/Pests/Phloeosinus-cupressi/. Consulted on 15 February 2021.

Flechtmann, C. A., & Atkinson, T. H. (2018). Oldest record of *Cyrtogenius luteus* (Blandford) (Coleoptera : Curculionidae: Scolytinae) from South America with notes on its distribution in Brazil. *Insecta Mundi* 0645: 1 – 4.

Foit, J., Kasak, J., Majek, T., Knizek, M., Hoch, G., Steyrer, G. (2017). First observations on the breeding ecology of invasive *Dryocoetes himalayensis* Strohmeyer, 1908 (Coleoptera: Curculionidae: Scolytinae) in its introduced range in Europe–Short communication. *Journal of Forest Science*, *63*(6), 290-292.

Gaaliche, B., Abdelaali, N. B., Mouttet, R., Halima-Kamel, M. B., & Hajlaoui, M. R. (2018). Nouveau signalement de *Hypocryphalus scabricollis* (Eichhoff, 1878) en Tunisie, un ravageur émergent sur figuier (*Ficus carica* L.). *EPPO Bulletin*, *48*(1), 164–166.

Galko, J., Dzurenko, M., Ranger, C. M., Kulfan, J., Kula, E., Nikolov, C., Zubrik, M., Zach, P. (2019). Distribution, Habitat Preference, and Management of the Invasive Ambrosia Beetle *Xylosandrus germanus* (Coleoptera: Curculionidae, Scolytinae) in European Forests with an Emphasis on the West Carpathians. *Forests*, *10*(1), 10; https://doi.org/10.3390/f10010010.

Garonna, A. P., Dole, S. A., Saracino, A., Mazzoleni, S., & Cristinzio, G. 2012. First record of the black twig borer *Xylosandrus compactus* (Eichhoff) (Coleoptera: Curculionidae, Scolytinae) from Europe. *Zootaxa*, 3251: 64-68.

Goldarazena, A., D. E. Bright, S. M. Hishinuma, S. Lopez, and S. J. Seybold. 2014. First record of *Pityophthorus solus* (Blackman, 1928) in Europe. *EPPO Bulletin* 44:65–69.

Gómez DF, Rabaglia RJ, Fairbanks KE, Hulcr J. 2018. North American Xyleborini north of Mexico: a review and key to genera and species (Coleoptera, Curculionidae, Scolytinae). *ZooKeys*, 768, 19.

Gómez, D., Martinez, G., & Beaver, R. A. (2012). First record of *Cyrtogenius luteus* (Blandford) (Coleoptera: Curculionidae: Scolytinae) in the Americas and its distribution in Uruguay. *The Coleopterists Bulletin*, *66*(4), 362–365.

Gómez, D., Suárez, M., & Martínez, G. (2017). *Amasa truncata* (Erichson) (Coleoptera: Curculionidae: Scolytinae): A new exotic ambrosia beetle in Uruguay. *The Coleopterists Bulletin*, *71*(4), 825–827.

Gómez, D.F.; Skelton, J.; Steininger, M.S.; Stouthamer, R.; Rugman-Jones, P.; Sittichaya, W.; Rabaglia, R.J.; Hulcr, J. 2018b. Species Delineation Within the *Euwallacea fornicatus* (Coleoptera: Curculionidae) Complex Revealed by Morphometric and Phylogenetic Analyses. *Insect Systematics and Diversity*, 2(6). doi:10.1093/isd/ixy018

Table S-1c. References


González, R., & Campos, M. (1994). A preliminary study of the effect of attacks by Phloetribus scarabaeoides (Bern.)(Coleoptera: Scolytidae) on the productiv ity of the olive tree (Olea europaea L.). *Mitteilungen der Schweizerischen Entomologischen Gesellschaft*, *67*(1-2), 67-75.

Grégoire, J.-C. 1988. The Greater European Spruce Beetle, *Dendroctonus micans* (Kug.). Pp. 455–478 *in* A.A. Berryman (Ed.), *Population Dynamics of Forest Insects,* Plenum, New-York, 604 pp.

Haack R. A. 2001. Intercepted Scolytidae (Coleoptera) at U.S. ports of entry: 1985–2000. *Integrated Pest Management Reviews,* 6: 253 – 282.

Haack R. A. 2006. Exotic bark-and wood-boring Coleoptera in the United States: recent establishments and interceptions. *Canadian Journal of Forest Research*, 36 (2): 269–288.

Hedgren, P. O. (2004), The bark beetle *Pityogenes chalcographus* (L.) (Scolytidae) in living trees: reproductive success, tree mortality and interaction with *Ips typographus*. *Journal of Applied Entomology*, 128: 161-166. doi:10.1046/j.1439-0418.2003.00809.x

Hoebeke ER, Rabaglia RJ, Knižek M, Weaver JS (2018) First records of *Cyclorhipidion fukiense* (Eggers) (Coleoptera: Curculionidae: Scolytinae: Xyleborini), an ambrosia beetle native to Asia, in North America. *Zootaxa* 4394: 243 – 250.

Holusa, J., Foit, J., Knizek, M., Schovankova, J., Lukasova, K., Vanicka, H., Trombik, J., Kula, E. (2019). The bark beetles *Orthotomicus laricis* and *Orthotomicus longicollis* are not pests in Central Europe: a case study from the Czech Republic. *Bulletin of Insectology*, *72*(2), 253-260.

Hughes, M. A., Riggins, J. J., Koch, F. H., Cognato, A. I., Anderson, C., Formby, J. P., Dreaden, T., Ploetz, R., Smith, J. A. (2017). No rest for the laurels: symbiotic invaders cause unprecedented damage to southern USA forests. *Biological Invasions*, *19*(7), 2143–2157.

Hulcr, J., Skelton, J., Johnson, A. J., Li, Y., & Jusino, M. A. (2018). Invasion of an inconspicuous ambrosia beetle and fungus may alter wood decay in Southeastern North America (No. e27334v1). *PeerJ* Preprints.

Inward, D. J. (2020). Three new species of ambrosia beetles established in Great Britain illustrate unresolved risks from imported wood. *Journal of Pest Science*, *93*(1), 117-126.

Israelson G (1990) A key to the Macaronesian Hypoborini, with description of two new species (Coleoptera, Scolytidae). *Bocagiana* 137: 1 – 11.

Jaramillo, J., Torto, B., Mwenda, D., Troeger, A., Borgemeister, C., Poehling, H. M., & Francke, W. (2013). Coffee berry borer joins bark beetles in coffee klatch. *PLoS One*, *8*(9), e74277.

Johnson, A. J., Knižek, M., Atkinson, T. H., Jordal, B. H., Ploetz, R. C., & Hulcr, J. (2017). Resolution of a Global Mango and Fig Pest Identity Crisis. Insect Systematics and Diversity, 1(2). doi:10.1093/isd/ixx010

Jordal, B. H., & Kirkendall, L. R. (1998). Ecological relationships of a guild of tropical beetles breeding in *Cecropia* petioles in Costa Rica. *Journal of Tropical Ecology*, 14(2), 153–176.

Kambestad, M., Kirkendall, L. R., Knutsen, I. L., & Jordal, B. H. (2017). Cryptic and pseudo-cryptic diversity in the world's most common bark beetle – *Hypothenemus eruditus*. *Organisms Diversity & Evolution*, *17*(3), 633–652.

Kirkendall, L. R. (2018). Invasive bark beetles (Coleoptera, Curculionidae, Scolytinae) in Chile and Argentina, including two species new for South America, and the correct identity of the Orthotomicus species in Chile and Argentina. *Diversity*, 10(2), 40.

Kirkendall L. R., Faccoli M. 2010. Bark beetles and pinhole borers (Curculionidae, Scolytinae, Platypodinae) alien to Europe. *Zookeys,* 56: 227 – 251.

Kirkendall LR, Biedermann PHW, Jordal BH. 2015. Evolution and Diversity of Bark and Ambrosia Beetles. Pp. 85–156 in *Bark Beetles, Biology and Ecology of Native and Invasive Species*, 1st Edition, Vega F and Hofstetter R (eds), Academic Press

Kirkendall, L. R., Dal Cortivo, M., & Gatti, E. (2008). First record of the ambrosia beetle, Monarthrum mali (Curculionidae, Scolytinae) in Europe. *Journal of Pest Science*, *81*(3), 175-178.

Kolařík, M., Kostovčík, M., & Pažoutová, S. (2007). Host range and diversity of the genus *Geosmithia* (Ascomycota: Hypocreales) living in association with bark beetles in the Mediterranean area. *Mycological Research*, *111*(11), 1298-1310.

Krivets, S.A., Bisirova, E.M., Kerchev, I.A. *et al.* 2015. Transformation of taiga ecosystems in the Western Siberian invasion focus of four-eyed fir bark beetle *Polygraphus proximus* Blandford (Coleoptera: Curculionidae, Scolytinae). *Russ J Biol Invasions* **6,** 94–108.

Kumbaşli, M., Hizal, E., Acer, S., Arslangündoğdu, Z., & Aday Kaya, A. G. (2018). First record of *Hylaste opacus* Erichson and *Crypturgus hispidulus* Thomson (Coleoptera; Curculionidae; Scolytinae) for the Turkish fauna. *Applied Ecology and Environmental Research*, *16*(4), 4585-4591.

La Spina S, De Cannière C, Dekri A, Grégoire J-C (2013) Frost increases beech susceptibility to scolytine ambrosia beetles. *Agric For Entomol* 15:157 – 167.

Langstrom, B., Hellqvist, C., 1993. Induced and spontaneous attacks by pine shoot beetles on young Scots pine trees—tree mortality and beetle performance. *J. Appl. Entomol.* 115, 25–36.

Lantschner, M.V., Corley, J.C. and Liebhold, A.M. (2020), Drivers of global Scolytinae invasion patterns. *Ecol Appl.* 30: e02103. doi:10.1002/eap.2103

Lee JC, Aguayo I, Aslin R, Durham G, Hamud SM, Moltzan BD, Munson AS, Negron JF, Peterson T, Ragenovich IR, Witcosky JJ, Seybold SJ (2009) Co-occurrence of the invasive banded and European elm bark beetles (Coleoptera: Scolytidae) in North America. *Annals of the Entomological Society of America*, 102 (3), 426–436.

Liu Z., Xu, B., Miao, Z., Sun, J. 2013. The Pheromone Frontalin and its Dual Function in the Invasive Bark Beetle *Dendroctonus valens*. *Chemical Senses*, 38 (6), 485–495. https://doi.org/10.1093/chemse/bjt019.

Mandelshtam, M.Y., Yakushkin, E.A. & Petrov, A.V. Oriental Ambrosia Beetles (Coleoptera: Curculionidae: Scolytinae): New Inhabitants of Primorsky Krai in Russia. *Russ J Biol Invasions* **9,** 355–365 (2018). https://doi-org.ezproxy.ulb.ac.be/10.1134/S2075111718040082

### Table S-1c. References


Mendel Z, Protasov A, Sharon M, Zveibil A, Ben Yehuda S, O'Donnell K, Rabaglia R, Wysoki M, Freeman S (2012) An Asian ambrosia beetle *Euwallacea fornicatus* and its novel symbiotic fungus *Fusarium* sp. pose a serious threat to the Israeli avocado industry. *Phytoparasitica* 40:235 – 238

Mendel, Z. (1984). Life history of *Phloeosinus armatus* Reiter and *P. aubei* Perris (Coleoptera: Scolytidae) in Israel. *Phytoparasitica*, *12*(2), 89-97.

Mendel, Z. (1987). Major pests of man-made forests in Israel: origin, biology, damage and control. *Phytoparasitica*, *15*(2), 131-137.

Mendel, Z., Assael, F., Saphir, N., Zehavi, A., Nestel, D., & Schiller, G. (1997). Seedling mortality in regeneration of Aleppo pine following fire and attack by the scale insect Matsucoccus josephi. *International Journal of Wildland Fire*, *7*(4), 327-333.

Menocal, O., Cruz, L. F., Kendra, P. E., Crane, J. H., Ploetz, R. C., & Carrillo, D. (2017). Rearing *Xyleborus volvulus* (Coleoptera: Curculionidae) on media containing sawdust from avocado or silkbay, with or without Raffaelea lauricola (Ophiostomatales: Ophiostomataceae). *Environmental entomology*, *46*(6), 1275-1283.

Mifsud, D., & Knizek, M. (2009). The bark beetles (Coleoptera: Scolytidae) of the Maltese islands (central Mediterranean). B ulletin of the Entomological Society of Malta, 2, 25–52.

Montecchio L., Faccoli M. 2014. First record of thousand cankers disease *Geosmithia morbida* and walnut twig beetle *Pityophthorus juglandis* on *Juglans nigra* in Europe. *Plant Disease*, 98 (5): 696-696.

Moraal, L. G. (2010). Infestations of the cypress bark beetles *Phloeosinus rudis*, *P. bicolor* and *P. thujae* in The Netherlands (Coleoptera: Curculionidae: Scolytinae). *entomologische berichten*, *70*(4), 140-145.

Moucheron, B., Dahan, L., Delbol, M., Ignace, D., Limbourg, P., Raemdonck, H., Drulmont, A. 2019. *Phloeosinus rudis* Blandford, 1894, scolyte invasif et nouveau pour la faune belge. *Lambillionea*, CXIX 1: 25-33

Nakagawa, M., Itioka, T., Momose, K., Yumoto, T., Komai, F., Morimoto, K., Jordal, B.H., Kato, M., Kaliang, H., Hamid, A.A., Inoue, T., Nakashizuka, T. (2003). Resource use of insect seed predators during general flowering and seeding events in a Bornean dipterocarp rain forest. *Bulletin of entomological research*, *93*(5), 455–466.

Neumann, F. G. (1987). Introduced bark beetles on exotic trees in Australia with special reference to infestations of Ips grandicollis in pine plantations. *Australian Forestry*, *50*(3), 166-178.

O'Donnell, K., Libeskind-Hadas, R., Hulcr, J., Bateman, C., Kasson, M. T., Ploetz, R. C., Carrillo, D., Campbell, A., Duncan, R. E., Liyanage, P. N. H., Eskalen, A., Lynch, S.C., Geiser, D. M., Freeman, S., Mendel, Z., Sharon, M., Aoki, T., Cossé, A. C., Rooney, A. (2016). Invasive Asian *Fusarium – Euwallacea* ambrosia beetle mutualists pose a serious threat to forests, urban landscapes and the avocado industry. *Phytoparasitica*, *44*(4), 435–442.

Okins KE, Thomas MC (2010) A new North American record for *Xyleborinus andrewesi* (Coleoptera: Curculionidae: Scolytinae). *Florida Entomologist* 93(1): 133–134.

Okins, K. E., & Thomas, M. C. (2009). Another Asian Ambrosia Bettle Established in Florida (Coleoptera: Curculionidae: Scolytinae: Xyleborini). *Florida Department of Agriculture and Consumer Services, Division of Plant Industry Charles H. Bronson, Commissioner of Agriculture, FL, USA*.

Paap, T., De Beer, Z. W., Migliorini, D., Nel, W. J., & Wingfield, M. J. (2018). The polyphagous shot hole borer (PSHB) and its fungal symbiont *Fusarium euwallaceae*: a new invasion in South Africa. *Australasian Plant Pathology*, *47*(2), 231–237.

Pennacchio, F., Faggi, M., Gatti, E., Caronni, F., Colombo, M., Roversi, P. F. (2004). First record of *Phloeotribus liminaris* (Harris) in Europe (Coleoptera Scolytidae). *Redia*, *87*, 85-89.

Rabaglia RJ, Dole SA, Cognato AI (2006) Review of American Xyleborina (Coleoptera: Curculionidae: Scolytinae) occurring north of Mexico, an illustrated key. *Ann Entomol Soc Am* 99:1034 – 1056

Rabaglia RJ, Smith SL, Rugman-Jones P, DiGirolomo M, Ewing C, Eskalen A (2020) Establishment of a non-native xyleborine ambrosia beetle, *Xyleborus monographus* (Fabricius) (Coleoptera: Curculionidae: Scolytinae), new to North America in California. Zootaxa 4786: 269–276. https://doi.org/10.11646/zootaxa.4786.2.8

Rabaglia RJ, Vandenberg NJ, Acciavatti, RE. 2009. First records of *Anisandrus maiche* Stark (Coleoptera: Curculionidae: Scolytinae) in North America. *Zootaxa* 2137: 23 – 28.

Rabaglia, R. J., M. Knızek & W. Johnson. 2010. First records of *Xyleborinus octiesdentatus* (Murayama) (Coleoptera, Curculionidae, Scolytinae) from North America. In: Cognato, A. I.; Knızek, M. (eds.) Sixty years of discovering scolytine and platypodine diversity: A tribute to Stephen L. Wood. *ZooKeys* 56:219 – 226.

Reay, S. D., & Walsh, P. J. (2002). The incidence of seedling attack and mortality by *Hylastes ater* (Coleoptera: Scolytidae) in second rotation Pinus radiata forests in the central North Island, New Zealand. *New Zealand Journal of Forestry*, *47*(2), 19-23.

Rodriguez, M., Delibes, M., & Fedriani, J. M. (2014). Hierarchical levels of seed predation variation by introduced beetles on an endemic Mediterranean palm. *PloS one*, 9(10), e109867.

Sanguansub, S., Goto, H., & Kamata, N. (2012). Guild structure of ambrosia beetles attacking a deciduous oak tree *Quercus serrata* in relation to wood oldness and seasonality in three locations in the Central Japan. *Entomological science*, *15*(1), 42-55.

Schedl, K. E. (1957). Bark-and timber-beetles from South Africa. *Annals and Magazine of Natural History*, *10*(110), 149-159.

Schedl, K. E. (1965). South African bark and timber beetles. *Journal of the Entomological Society of Southern Africa*, *28*(1), 110-116.

Seybold, S. J., Penrose, R. L., & Graves, A. D. (2016). Invasive bark and ambrosia beetles in California Mediterranean forest ecosystems. In *Insects and diseases of Mediterranean forest systems* (pp. 583–662). Springer International Publishing Switzerland.

Seybold, Steve; Dallara, Paul L.; Nelson, Lori; Graves, Andrew D.; Hishinuma, Stacy M; Gries, Regine. 2015. Patent: Methods of Monitoring and Controlling the Walnut Twig Beetle, Pityophthorus Juglandis. https://patents.google.com/patent/US9137990B2/en?oq=13548319.

### **Table S-1c.** References


Shapovalov, M. I., Zamotajlov, A. S., Saprykin, M. A. 2010. Coleopterous insects (Insecta, Coleoptera) of Republic of Adygheya (annotated catalogue of species) (Fauna conspecta of Adygheya. № 1)/Edited by A. S. Zamotajlov and N. B. Nikitsky. – Maykop: Adyghei State University Publishers, 404 pp.

Six, D. L., De Beer, Z. W., Beaver, R. A., Visser, L., & Wingfield, M. J. (2005). Exotic invasive elm bark beetle, *Scolytus kirschii*, detected in South Africa: research in action. *South African journal of science*, *101* (5–6), 229–232.

Björklund, N., & Boberg, J. (2017). Rapid Pest Risk Analysis *Xyleborinus attenuatus*. https://pub.epsilon.slu.se/id/eprint/15311. Consulted on 17 April 2020.

Smith, S. M., & Hulcr, J. (2015). *Scolytus* and other economically important bark and ambrosia beetles. Pp. 495–531 in *Bark Beetles, Biology and Ecology of Native and Invasive Species*, 1st Edition, Vega F and Hofstetter R (eds), Academic Press.

Smith, S. M., Beaver, R. A., & Cognato, A. I. (2020). A monograph of the Xyleborini (Coleoptera, Curculionidae, Scolytinae) of the Indochinese Peninsula (except Malaysia) and China. *ZooKeys*, *983*, 1-442. doi: 10.3897/zookeys.983.52630

Smith, S. M., Gomez, D. F., Beaver, R.A., Hulcr, J., Cognato, A. I. (2019). Reassessment of the Species in the *Euwallacea fornicatus* (Coleoptera: Curculionidae: Scolytinae) Complex after the Rediscovery of the “Lost” Type Specimen. *Insects*, *10*(9), 261.

Smith, S. M.; Beaver, R.A.; Cognato, A.I. 2017. The ambrosia beetle *Ambrosiophilus peregrinus* Smith and Cognato, introduced to the USA, is *Ambrosiophilus nodulosus* (Eggers), new combination (Coleoptera: Curculionidae: Scolytinae). *Coleopterists Bull.* 71(3): 552–553.

Smith, S.M. , Cognato, A.I. 2015. *Ambrosiophilus peregrinus* Smith and Cognato, New Species (Coleoptera: Curculionidae: Scolytinae), an Exotic Ambrosia Beetle Discovered in Georgia, USA. *Coleopterists Bull.* 69(2): 213–220.

Sousa, W. P., Quek, S. P., & Mitchell, B. J. (2003). Regeneration of *Rhizophora mangle* in a Caribbean mangrove forest: interacting effects of canopy disturbance and a stem-boring beetle. *Oecologia*, *137*(3), 436-445.

Terekhova, V. V., & Skrylnik, Y. Y. (2012). Biological peculiarities of the alien for Europe Anisandrus maiche Stark (Coleoptera: Curculionidae: Scolytinae) bark beetle in Ukraine. *Russian Journal of Biological Invasions*, *3*(2), 139-144.

Tribe, G. D. (1992). Colonisation sites on *Pinus radiata* logs of the bark beetles, *Orthotomicus erosus*, *Hylastes angustatus* and *Hylurgus ligniperda* (Coleoptera: Scolytidae). *Journal of the Entomological Society of Southern Africa*, *55*(1), 77–84.

Vega, F. E., Infante, F., & Johnson, A. J. (2015). The genus *Hypothenemus*, with emphasis on *H. hampei*, the coffee berry borer. Pp. 427-494 in *Bark Beetles, Biology and Ecology of Native and Invasive Species*, 1st Edition, Vega F and Hofstetter R (eds), Academic Press.

Wood, S. L. 1977. Introduced and exported American Scolytidae (Coleoptera). *The Great Basin Naturalist*, 67-74.

Wood, S. L. 1982. The bark and ambrosia beetles of North and Central America (Coleoptera: Scolytidae), a taxonomic monograph. *Great Basin Nat. Mem.* 6:1-1356.

Wood, S. L., 2007: Bark and ambrosia beetles of South America (Coleoptera: Scolytidae). Bark and ambrosia beetles of South America. Monte L. Bean Sci. Mus., Provo, Utah: 1–900.

Wood, S. L., Bright Jr, D.E. 1992. A catalog of Scolytidae and Platypodidae (Coleoptera), Part 2. Taxonomic Index (Volumes A, B). *Great Basin Nat. Mem.* 13: 1-1553.

Yan, Z., Sun, J., Don, O., & Zhang, Z. (2005). The red turpentine beetle, *Dendroctonus valens* LeConte (Scolytidae): an exotic invasive pest of pine in China. *Biodiversity & Conservation*, *14*(7), 1735–1760.

Zanuncio, J. C., Sossai, M. F., Flechtmann, C. A. H., Zanuncio, T. V., Guimarães, E. M., & Espindula, M. C. (2005). Plants of an Eucalyptus clone damaged by Scolytidae and Platypodidae (Coleoptera). *Pesquisa Agropecuária Brasileira*, *40*(5), 513-515.

Zeiri, A., Ahmed, M., Cuthbertson, A., & Braham, M. (2018). Monitoring the attack incidences and damage caused by the almond bark beetle, *Scolytus amygdali*, in almond orchards. *Insects*, *9*(1), 1.
