## Supplemental table S2 for "Cosmopolitan Scolytinae: strong common drivers but too many singularities for accurate prediction"

**Table S-2.** Landmasses (islands and continents).

| Continent, country, or island | ISO alpha-3 codes[1] | TDWG[2] | Additional [3] |
|---|---|---|---|
| Africa | | | AFR |
| Andaman & Nicobar Islands | | | AND |
| Antigua and Barbuda | ATG | | |
| Asia | | | ASI |
| Australia | AUS | | |
| Azores | | AZO | |
| Bahamas | BHS | | |
| Barbados | BRB | | |
| Bermuda | BMU | | |
| Bismarck islands | | BIS | |
| Bonin islands (JPN) | | | BON |
| Borneo (Indonesia) | | BOR | |
| Canary Islands | | CNY | |
| Cabo Verde Islands | CPV | | |
| Caroline Islands | | CRL | |
| Celebes Islands | | | CEL |
| Central American mainland | | | CAM |
| Christmas islands | CXR | | |
| Cocos islands | CCK | | |
| Comoros | COM | | |
| Cook Islands | COK | | |
| Corsica | | COR | |
| Cuba | CUB | | |
| Cyprus | CYP | | |
| Dominica | DMA | | |
| Europe | EUR | | |
| Fernando Poo | | | FER |
| Fiji | FJI | | |
| Galapagos | | GAL | |
| Great Britain | GBR | | |
| Gilbert islands | | GIL | |
| Grenada | GRD | | |
| Guadeloupe | GLP | | |
| Guam | GUM | | |
| Hainan | | | HAI |
| Haiti + Dominican Rep (Hispaniola island) | | | HIS |
| Hawaii (US) | | HAW | |
| Henderson Island | | | HEN |
| Hokkaido (JPN) | | | HOK |
| Honshu (JPN) | | | HON |
| Indonesia | IDN | | |
| Ireland | IRL | | |
| Jamaica | JAM | | |
| Japan | JPN | | |
| Java (IDN) | | JAW | |
| Kiribati | | | KIR |
| Kuril Islands | | KUR | |
| La Vega | | | LAV |
| Luzon (PHL) | | | LUZ |
| Madagascar | MDG | | |
| Madeira | | MDR | |
| Malta | MLT | | |
| Marianna islands | | | MAR |
| Marquesas | | MRQ | |
| Marshall | MHL | | |
| Martinique | MTQ | | |
| Mauritius | MUS | | |
| Mentawai (IDN) | | | MEN |

[1] https://www.iso.org/obp/ui/#search. Locations not in the list are in brackets
[2] International Working Group on Taxonomic Databases For Plant Sciences (TDWG) https://github.com/tdwg/wgsrpd
[3] Landmasses not listed in the two previous databases

**Table S-2.** Landmasses (islands and continents).

| Continent, country, or island | ISO alpha-3 codes[1] | TDWG[2] | Additional [3] |
|---|---|---|---|
| Micronesia | FSM | | |
| Mindanao (PHL) | | | MIN |
| Monserrat | MSR | | |
| Nevis | | | NEV |
| New Britain | | | NBR |
| New Caledonia | NCL | | |
| New Guinea | | NWG | |
| New Hebrides | | VAN | |
| New Zealand | NZL | | |
| Niue Island | | NUE | |
| North American mainland | | | NAM |
| Okinawa (Japan) | | | OKI |
| Palau | PLW | | |
| Papua New Guinea | PNG | | |
| Philippines | PHL | | |
| Puerto Rico | PRI | | |
| Réunion | REU | | |
| Saint Helena | SHN | | |
| Saint Kitts and Nevis | KNA | | |
| Saint Vincent | VCT | | |
| Sakhalin (RU) | | SAK | |
| Samoa | WSM | | |
| Santa Lucia | LCA | | |
| Sao Tomé | STP | | |
| Sardinia (ITA) | | SAR | |
| Seychelles | SYC | | |
| Sicily (ITA) | | SIC | |
| Society Island | | SCI | |
| Solomon Islands | SLB | | |
| South American mainland | | | SAM |
| Sri lanka | LKA | | |
| Sumatra (IDN) | | SUM | |
| Tahiti (FRA) | | | TAH |
| Taïwan | TWN | | |
| Tasmania (AUS) | | TAS | |
| Tokelau | TKL | | |
| Tonga | TON | | |
| Trinidad & Tobago | TTO | | |
| Tuamotu Islands | | TUA | |
| Virgin Islands (GB + US) | | | VIL |

Conventions:

- Acronyms: ISO alpha-3, or TDWG codes, or created on purpose (e.g. for whole continents, or for islands part of a wider country).
- Malaysia: by default: on continental Asia, except if explicitely on Borneo island (Sarawak).
- Haiti + Dominican Republic = Hispaniola island (HIS).
- Middle East: Asia.
- EUR: Europe as a whole (EU + other European countries)
