## Supplemental table S3 for "Cosmopolitan Scolytinae: strong common drivers but too many singularities for accurate prediction"

**Table S-3.** Well classified and mis-classified species identified by the Factorial Discriminant Analysis (FDA) of the three categories of impact by the 163 beetle species studied

| Species | *A priori* classification [1] | *A posteriori* classification [2] | Comment |
|---|---|---|---|
| *Ambrosiodmus lewisi* (Blandford) | 0 | 0 | well classified |
| *Ambrosiodmus minor* (Stebbing) | 0 | 0 | well classified |
| *Ambrosiodmus obliquus* (LeConte) | 0 | 0 | well classified |
| *Ambrosiophilus nodulosus* (Eggers) | 0 | 0 | well classified |
| *Anisandrus maiche* (Kurentzov) | 0 | 0 | well classified |
| *Aphanarthrum affine* Wollaston | 0 | 0 | well classified |
| *Aphanarthrum* (*Coleobothrus*) *alluaudi* Peyerimhoff | 0 | 0 | well classified |
| *Aphanarthrum bicolor* Wollaston | 0 | 0 | well classified |
| *Aphanarthrum mairei* Peyerimhoff | 0 | 0 | well classified |
| *Aphanarthrum piscatorium* Wollaston | 0 | 0 | well classified |
| *Coccotrypes aciculatus* Schedl | 0 | 0 | well classified |
| *Coccotrypes advena* Blandford | 0 | 0 | well classified |
| *Coccotrypes cyperi* (Beeson) | 0 | 0 | well classified |
| *Coccotrypes distinctus* (Motschulsky) | 0 | 0 | well classified |
| *Coccotrypes robustus* Eichhoff | 0 | 0 | well classified |
| *Coccotrypes rutschuruensis* Eggers | 0 | 0 | well classified |
| *Coccotrypes vulgaris* (Eggers) | 0 | 0 | well classified |
| *Cryphalus wapleri* Eichhoff | 0 | 0 | well classified |
| *Cryptocarenus heveae* (Hagedorn) | 0 | 0 | well classified |
| *Cryphalus pallidus* Eichhoff | 0 | 0 | well classified |
| *Crypturgus cylindricollis* Eggers | 0 | 0 | well classified |
| *Crypturgus numidicus* Ferrari | 0 | 0 | well classified |
| *Crypturgus pusillus* (Gyllenhal) | 0 | 0 | well classified |
| *Cyclorhipidion fukiense* (Eggers) | 0 | 0 | well classified |
| *Cyrtogenius luteus* (Blanford) | 0 | 0 | well classified |
| *Dactylotrypes longicollis* (Wollaston) | 0 | 0 | well classified |
| *Dryocoetoides cristatus* (Fabricius) | 0 | 0 | well classified |
| *Dryocoetes himalayensis* Strohmeyer | 0 | 0 | well classified |
| *Dryoxylon onoharaense* (Murayama) | 0 | 0 | well classified |
| *Eccoptopterus spinosus* (Olivier) | 0 | 0 | well classified |
| *Planiculus (Euwallacea) bicolor* (Blandford) | 0 | 0 | well classified |
| *Gnathotrichus materiarius* (Fitch) | 0 | 0 | well classified |
| *Hylastes linearis* Erichson | 0 | 0 | well classified |
| *Hylastes opacus* Erichson | 0 | 0 | well classified |
| *Hylastinus obscurus* (Marsham) | 0 | 0 | well classified |
| *Hylesinus toranio* (= *Bostrychus oleiperda*) (Danthione) | 0 | 0 | well classified |
| *Hylurgus ligniperda* (Fabricius) | 0 | 0 | well classified |
| *Hylurgus micklitzi* Wachtl | 0 | 0 | well classified |
| *Hylurgops palliatus* (Gyllenhal) | 0 | 0 | well classified |
| *Hypothenemus africanus* (Hopkins) | 0 | 0 | well classified |

[1] According to the literature (see Table S-3)

[2] According to our Factorial Discriminant Analysis (see section 2.5)

**Table S-3.** Well classified and mis-classified species identified by the Factorial Discriminant Analysis (FDA) of the three categories of impact by the 163 beetle species studied

| **Species** | ***A priori*** **classification** 1 | ***A posteriori*** **classification** 2 | **Comment** |
|---|---|---|---|
| *Hypothenemus brunneus* Hopkins | 0 | 0 | well classified |
| *Hypothenemus californicus* Hopkins | 0 | 0 | well classified |
| *Hypothenemus columbi* Hopkins | 0 | 0 | well classified |
| *Hypothenemus erectus* LeConte | 0 | 0 | well classified |
| *Hypothenemus javanus* (Eggers) | 0 | 0 | well classified |
| *Hypothenemus leprieuri* ( Perris) | 0 | 0 | well classified |
| *Hypothenemus pubescens* Hopkins | 0 | 0 | well classified |
| *Hypothenemus setosus* (Eichhoff) | 0 | 0 | well classified |
| *Hypocryphalus dilutus* (Eichhoff) | 0 | 0 | well classified |
| *Hypothenemus elephas* (Eichhoff) | 0 | 0 | well classified |
| *Ips calligraphus* (Germar) | 0 | 0 | well classified |
| *Orthotomicus laricis* (Fabricius) | 0 | 0 | well classified |
| *Orthotomicus proximus* (Eichhoff) | 0 | 0 | well classified |
| *Kissophagus hederae* (Schmidt) | 0 | 0 | well classified |
| *Liparthrum artemisiae* Wollaston | 0 | 0 | well classified |
| *Liparthrum bituberculatum* Wollaston | 0 | 0 | well classified |
| *Liparthrum curtum* Wollaston | 0 | 0 | well classified |
| *Liparthrum inarmatum* Wollaston | 0 | 0 | well classified |
| *Liparthrum mandibulare* Wollaston | 0 | 0 | well classified |
| *Liparthrum mori* (Aube) | 0 | 0 | well classified |
| *Microborus boops* Blandford | 0 | 0 | well classified |
| *Microperus quercicola* (Eggers) | 0 | 0 | well classified |
| *Microperus* (*Coptodryas*) *eucalypticus* (Schedl) | 0 | 0 | well classified |
| *Monarthrum mali* (Fitch) | 0 | 0 | well classified |
| *Orthotomicus angulatus* (Eichhoff) | 0 | 0 | well classified |
| *Orthotomicus caelatus* (Eichhoff) | 0 | 0 | well classified |
| *Pagiocerus frontalis* (Fabricius) | 0 | 0 | well classified |
| *Phloeosinus cupressi* Hopkins | 0 | 0 | well classified |
| *Pityophthorus solus* (Blackman) | 0 | 0 | well classified |
| *Pityogenes bidentatus* (Herbst) | 0 | 0 | well classified |
| *Polygraphus poligraphus* (Linnaeus) | 0 | 0 | well classified |
| *Polygraphus rufipennis* (Kirby) | 0 | 0 | well classified |
| *Premnobius ambitiosus* (Schaufuss) | 0 | 0 | well classified |
| *Premnobius cavipennis* Eichhoff | 0 | 0 | well classified |
| *Pseudohylesinus sericeus* (Mannerheim) | 0 | 0 | well classified |
| *Scolytus dimidiatus* Chapuis | 0 | 0 | well classified |
| *Scolytogenes jalapae (Letzner)* | 0 | 0 | well classified |
| *Scolytus amygdali* Guerin-Meneville | 0 | 0 | well classified |
| *Scolytus mali* (Bechstein) | 0 | 0 | well classified |
| *Scolytus sulcifrons* Rey | 0 | 0 | well classified |
| *Scolytoplatypus tycon* Blandford | 0 | 0 | well classified |
| *Hypothenemus plumeriae* (Nordlinger) | 0 | 0 | well classified |
| *Theoborus ricini* (Eggers) | 0 | 0 | well classified |
| *Thamnurgus characiae* Rosenhauer | 0 | 0 | well classified |
| *Truncaudum* (*Cyclorhipidion*) *agnatum* (Eggers) | 0 | 0 | well classified |
| *Xyleborinus attenuatus* (Blandford) | 0 | 0 | well classified |

**Table S-3.** Well classified and mis-classified species identified by the Factorial Discriminant Analysis (FDA) of the three categories of impact by the 163 beetle species studied

| Species | *A priori* classification [1] | *A posteriori* classification [2] | Comment |
|---|---|---|---|
| *Xyleborinus andrewesi* (Blandford) | 0 | 0 | well classified |
| *Xyleborinus artestriatus* (Eichhoff) | 0 | 0 | well classified |
| *Xyleborinus exiguus* (Walker) | 0 | 0 | well classified |
| *Xyleborinus gracilis* Eichhoff | 0 | 0 | well classified |
| *Xyleborinus octiesdentatus* (Murayama) | 0 | 0 | well classified |
| *Xyleborus africanus* Eggers | 0 | 0 | well classified |
| *Xyleborus atratus* Eichhoff | 0 | 0 | well classified |
| *Ambrosiophilus* (*Xyleborus*) *atratus* (Eichhoff) | 0 | 0 | well classified |
| *Xyleborus bispinatus* Eichhoff | 0 | 0 | well classified |
| *Cyclorhipidion bodoanum* (*Xyleborus californicus*) (Reitter) | 0 | 0 | well classified |
| *Xyleborus monographus* (Fabricius) | 0 | 0 | well classified |
| *Xyleborus pfeilii* (Ratzeburg) | 0 | 0 | well classified |
| *Cyclorhipidion pelliculosum* (Eichhoff) | 0 | 0 | well classified |
| *Xyleborus seriatus* Blandford | 0 | 0 | well classified |
| *Xyleborus spinulosus* Blandford | 0 | 0 | well classified |
| *Xylosandrus* (*Apoxyleborus*) *mancus* (Blandford) | 0 | 0 | well classified |
| *Xylosandrus amputatus* (Blandford) | 0 | 0 | well classified |
| *Cnestus* (*Xylosandrus*) *mutilatus* (Blandford | 0 | 0 | well classified |
| *Xylosandrus pseudosolidus* (Schedl) | 0 | 0 | well classified |
| *Xyleborus volvulus* (Fabricius) | 0 | 0 | well classified |
| *Xyloterinus politus* (Say) | 0 | 0 | well classified |
| *Ambrosiodmus compressus* (Lea) | 1 | 0 | mis-classified |
| *Ambrosiodmus rubricollis* (Eichhoff) | 1 | 0 | mis-classified |
| *Tomicus piniperda* (Linnaeus) | 1 | 0 | mis-classified |
| *Coccotrypes carpophagus* (Hornung) | 1 | 0 | mis-classified |
| *Hypocryphalus* (*Cryphalus*) *scabricollis* (Eichhoff) | 1 | 0 | mis-classified |
| *Hylastes angustatus* (Herbst) | 1 | 0 | mis-classified |
| *Hylastes ater* (Paykull) | 1 | 0 | mis-classified |
| *Hypothenemus birmanus* (Eichhoff) | 1 | 0 | mis-classified |
| *Hypothenemus obscurus* (Fabricius) | 1 | 0 | mis-classified |
| *Hypothenemus areccae* (Hornung) | 1 | 0 | mis-classified |
| *Hypoborus ficus* Erichson | 1 | 0 | mis-classified |
| *Hypocryphalus mangiferae* (Stebbing) | 1 | 0 | mis-classified |
| *Ips cembrae* (Herr) | 1 | 0 | mis-classified |
| *Orthotomicus erosus* (Wollaston) | 1 | 0 | mis-classified |
| *Ips grandicollis* (Eichhoff) | 1 | 0 | mis-classified |
| *Phloeotribus liminaris* (Harris) | 1 | 0 | mis-classified |
| *Phloeotribus scarabaeoides* (Bernard) | 1 | 0 | mis-classified |
| *Phloeosinus armatus* Reitter | 1 | 0 | mis-classified |
| *Phloeosinus rudis* Blandford | 1 | 0 | mis-classified |
| *Phloeosinus thujae* (Perris) | 1 | 0 | mis-classified |
| *Pityokteines curvidens* (Germar) | 1 | 0 | mis-classified |
| *Pityogenes calcaratus* (Eichhoff) | 1 | 0 | mis-classified |
| *Pityogenes chalcographus* (Linnaeus) | 1 | 0 | mis-classified |
| *Polygraphus proximus* Blandford | 1 | 0 | mis-classified |

**Table S-3.** Well classified and mis-classified species identified by the Factorial Discriminant Analysis (FDA) of the three categories of impact by the 163 beetle species studied

| Species | *A priori* classification [1] | *A posteriori* classification [2] | Comment |
|---|---|---|---|
| *Scolytus rugulosus* (Muller) | 1 | 0 | mis-classified |
| *Trypodendron domesticum* (Linnaeus) | 1 | 0 | mis-classified |
| *Anisandrus* (*Xyleborus*) *dispar* (Fabricius) | 1 | 0 | mis-classified |
| *Xyleborus perforans* (Wollaston) | 1 | 0 | mis-classified |
| *Amasa truncatus* (*Xyleborus truncatus*) (Erichson) | 1 | 0 | mis-classified |
| *Euwallacea* (*Xyleborus*) *piceus* (Motschulsky) | 1 | 1 | well classified |
| *Hypothenemus eruditus* Westwood | 1 | 1 | well classified |
| *Hypothenemus seriatus* (Eichhoff) | 1 | 1 | well classified |
| *Xylosandrus germanus* (Blandford) | 1 | 1 | well classified |
| *Xyleborus affinis* Eichhoff | 1 | 2 | mis-classified |
| *Euwallacea (Xyleborus) similis (Ferrari)* | 1 | 2 | mis-classified |
| *Coccotrypes dactyliperda* Fabricius | 2 | 0 | mis-classified |
| *Coccotrypes rhizophorae* (Hopkins) | 2 | 0 | mis-classified |
| *Dendroctonus micans* (Kugelann) | 2 | 0 | mis-classified |
| *Dendroctonus valens* LeConte | 2 | 0 | mis-classified |
| *Euwallacea kuroshio* Gomez and Hulcr | 2 | 0 | mis-classified |
| *Euwallacea perbrevis (Schedl, 1951)* | 2 | 0 | mis-classified |
| *Euwallacea fornicatus* (Eichhoff) | 2 | 0 | mis-classified |
| *Pityophthorus juglandis* Blackman | 2 | 0 | mis-classified |
| *Scolytus kirschi* Skahtzky | 2 | 0 | mis-classified |
| *Scolytus multistriatus* (Marsham) | 2 | 0 | mis-classified |
| *Scolytus schevyrewi* Semenov | 2 | 0 | mis-classified |
| *Hypothenemus hampei* (Ferrari) | 2 | 0 | mis-classified |
| *Xyleborus glabratus* Eichhoff | 2 | 0 | mis-classified |
| *Xyleborinus saxeseni* (Ratzeburg) | 2 | 0 | mis-classified |
| *Xylosandrus compactus* (Eichhoff) | 2 | 0 | mis-classified |
| *Hypothenemus crudiae* (Panzer) | 2 | 1 | mis-classified |
| *Xyleborus ferrugineus* (Fabricius) | 2 | 1 | mis-classified |
| *Euwallacea interjectus* (Blandford) | 2 | 1 | mis-classified |
| *Euwallacea validus* (Eichhoff) | 2 | 1 | mis-classified |
| *Xylosandrus crassiusculus* (Motschulsky) | 2 | 2 | well classified |
| *Xylosandrus morigerus* (Blandford) | 2 | 2 | well classified |
